# Base editing and nanoparticle transfection of airway cell types essential for treatment of cystic fibrosis

**DOI:** 10.1101/2025.11.06.687048

**Authors:** Erin W. Kavanagh, Anya T. Joynt, Audrey R. Pion, Alice C. Eastman, Alianna I. Parr, Katherine L. Starego, Manav Jain, Sydney R Shannon, Edwin Yoo, Gregory A. Newby, Stephany Y. Tzeng, Neeraj Sharma, Jordan J Green, Garry R Cutting

## Abstract

Cystic Fibrosis (CF) is a life-limiting genetic disorder caused by deleterious variants in the *CFTR* gene that results in altered mucous impairing the airway epithelia. Durable correction of these variants in airway cells remain a therapeutic challenge for ∼10% of individuals unresponsive to CFTR modulators. A common disease-causing *CFTR* splice site variant was corrected in primary CF airway cells using base editor RNAs. Single-cell RNA sequencing revealed a remarkable increase in detectable *CFTR* transcript in most CF airway epithelial cell types with notable enrichment of *CFTR*-expressing ionocytes and secretory goblet cells. Progenitor basal cell subtypes were edited but they decreased as a fraction of total cells and *CFTR* expressing cells compared to unedited cells. CRISPR base editors delivered by polymeric nanoparticles (PNPs) facilitated functional rescue of CFTR to clinically meaningful levels in immortalized and primary airway cells. PNPs delivered reporter encoding RNA to progenitor airway cells in fully differentiated airway cultures. Vitronectin was a major component of the PNP corona that formed in vivo, but pre-incubation with vitronectin did not enhance delivery. Together, these findings validate a scalable, non-viral platform with significant translational promise for treating CF and other respiratory diseases involving respiratory epithelial cell dysfunction.

## Introduction

Current treatments for cystic fibrosis (CF) primarily focus on modulator drug therapies designed to correct malfunctioning CF transmembrane conductance regulator (CFTR) protein.[1] However, these modulators are ineffective for individuals with CF carrying variants that do not synthesize CFTR protein. CRISPR/Cas9-mediated base editing can correct DNA variants to allow production of sufficient protein to achieve clinically relevant levels of CFTR function.[2–7] Among these is the canonical splice variant c.2988+1G>A (legacy 3120+1G>A), the most common CF-causing variant in native Africans.[8,9] The variant ablates splicing of exon 18 leading to a frameshift, introduction of premature termination codons causing transcript instability due to mRNA degradation and loss of protein synthesis.[10] We have demonstrated that electroporation of a modified CRISPR-Cas9 adenine base editor (ABE) to primary human bronchial epithelial (HBE) cells carrying 3120+1G>A achieved 75% genome editing resulting in ∼40% wild type (WT) CFTR function.[2] These are encouraging results; however, successful translation to individuals affected with CF requires evidence that editing systems correct epithelial cell types where CFTR plays a key role in ion and fluid transport. In conjunction, delivery vehicles must be evaluated for efficient transfer of components that effect editing of airway epithelial cells and recovery of CFTR function.

Characterization of the transcriptome of individual cells by single cell RNA sequencing (scRNA-seq) has deeply informed our understanding of cellular composition of surface airway epithelial cells in various locations in the bronchial tree. This method enabled the discovery of ionocytes, a rare cell type that expresses the highest level of *CFTR* mRNA [11,12] and is a key mediator of airway surface liquid (ASL) volume, pH and ionic content.[13–16] Despite this critical role, ionocytes account for only a small fraction of epithelial cells (1-2%) in the airways of mice, ferrets and humans. Although secretory epithelial cells individually express *CFTR* at lower level than ionocytes, their commonness (∼20-40%) accounts for 40-60% of *CFTR* transcripts in large and small airways.[17,18] Basal cells, the progenitor cell type that is responsible for surface epithelial cell replenishment [19] also individually express modest levels of *CFTR* mRNA, but they account for 15% to 50% of airway cells directly assayed from human lungs.[17,18] These studies indicate that correction of CFTR, particularly in the small airways where CF lung pathology is most evident, should broadly encompass most subsets of secretory and basal cells in order to maximize its therapeutic effect. In addition, the rare ionocytes should be edited as they are key mediators of ASL.

Comparison of freshly obtained healthy and CF epithelial cells reveals relatively similar distributions however, CF samples have an increase in subsets of ciliated and secretory cells and a decrease in cycling basal cells.[17] When *CFTR* is assessed, airway cells from healthy and CF individuals have similar distributions except for ionocytes that exhibit two-fold increases in *CFTR* positive (+) cells and overall fraction of *CFTR* expression.[17] While studies of fresh samples of the airways is desirable, they are challenging to obtain and transport, especially from individuals carrying less common *CFTR* genotypes that are more likely to be selected for gene correction treatments. Expansion and differentiation of airway epithelial cells obtained from bronchial lavage or nasal turbinate brushing using rho kinase inhibitors (ROCK) and air-liquid interface (ALI) provide a flexible solution to studying *CFTR* variants and their response to therapy. All major cell types observed in fresh samples of CF and non-CF subjects are found in cultured cells with some differences in select subsets.[17,20,21] Furthermore, scRNA-seq of cultured nasal cells reveals that the distribution of *CFTR* expressing cells corresponds to the distribution in small airways, the primary site of disease in CF.[20,22] Overall, nasal and bronchial cells cultured in ALI provide a reasonable representation of the differentiation process *in vivo* and the cell types that should express CFTR for restoration of normal ion and fluid transport.

Of equal importance to knowing what cells need to have restored CFTR function is to be able to deliver therapeutic components to those cells. For the lungs, administration of aerosolized viral and non-viral delivery vehicles has been explored extensively, although barriers to efficient transfection of the CF lung still need to be surmounted.[23,24] Systemic delivery of non-viral vectors has shown utility for single gene disorders requiring correction in hepatocytes.[25] However, specialized formulations are required to deliver nanoparticles (NPs) to organs affected in CF such as the lungs and pancreas.[20,21] An alternative approach is to formulate nanomaterials inherently suited for lung transfection. One such class of materials are biodegradable polymeric NPs [26–28] that enable efficient delivery of reporter mRNAs to primary human airway cells *in vitro* and mouse lung epithelial cells *in vivo*.[29,30]

To determine which *CFTR* expressing cell types are edited and whether gene correction alters cell types upon differentiation, we performed scRNA-seq of primary cells after editing of the 3120+1G>A variant. To address transfer, we tested the efficiency of polymeric NPs encapsulating base editing machinery to edit in differentiated immortalized CFBE (Cystic Fibrosis Bronchial Epithelial) cells with an integrated *CFTR* expression minigene bearing the 3120+1 variant. This approach was extended to primary airway cells bearing 3120+1 prior to ALI differentiation then fully differentiated cells using polymeric NPs encapsulating GFP RNA. Together, these studies determined which cell types undergo gene editing and generate corrected *CFTR* mRNA transcripts, the effect of *CFTR* correction on respiratory cell types and ability of polymeric NPs to deliver editing cargo to differentiating human airway cells.

## Results

### Editing with different guide RNAs results in substantial increases in number of primary airway cells with detectable CFTR transcript

To ensure broad and highly efficient transfection, we used electroporation to transfect primary human nasal epithelial cells (HNE) from an individual bearing 3120+1G>A (c.2988+1G>A)/G480S (c.1438G>A) with ABE8e mRNA and one of two sg RNAs (sgRNA4long or sgRNA5long) with similar editing efficiency.[2] Edited and unedited cells were cultured in parallel using the same media and supplements. Single cell RNA-sequencing (scRNA-seq) was performed on cells differentiated for 21 days (**Figure 1A**). Transfection with RNA encoding green fluorescent protein (GFP) revealed that delivery approached 100% of HNE cells **(Figure 1B**). DNA editing of the target nucleotide in HNE cells was efficient, with 85% A>G conversion with sgRNA4long and 88% with sgRNA5long **(Figure 1C**). Conversion was calculated after taking into account the heterozygous status at the target site due to the presence of the other *CFTR* allele (G480S) and were comparable to previous reports.[2] The mean level of *CFTR* expression in cells having detectable *CFTR* transcript (i.e., *CFTR*+ cells) was similar between unedited (2.07) and edited cells (sg4long 1.69; sg5long 2.04) (**Figure 1D**). However, the number of cells with detectable *CFTR* transcript increased 5.2-fold between unedited (0.06) and sg4long (0.31) and 6.9-fold between unedited and sg5long (0.41).

**Figure 1:**
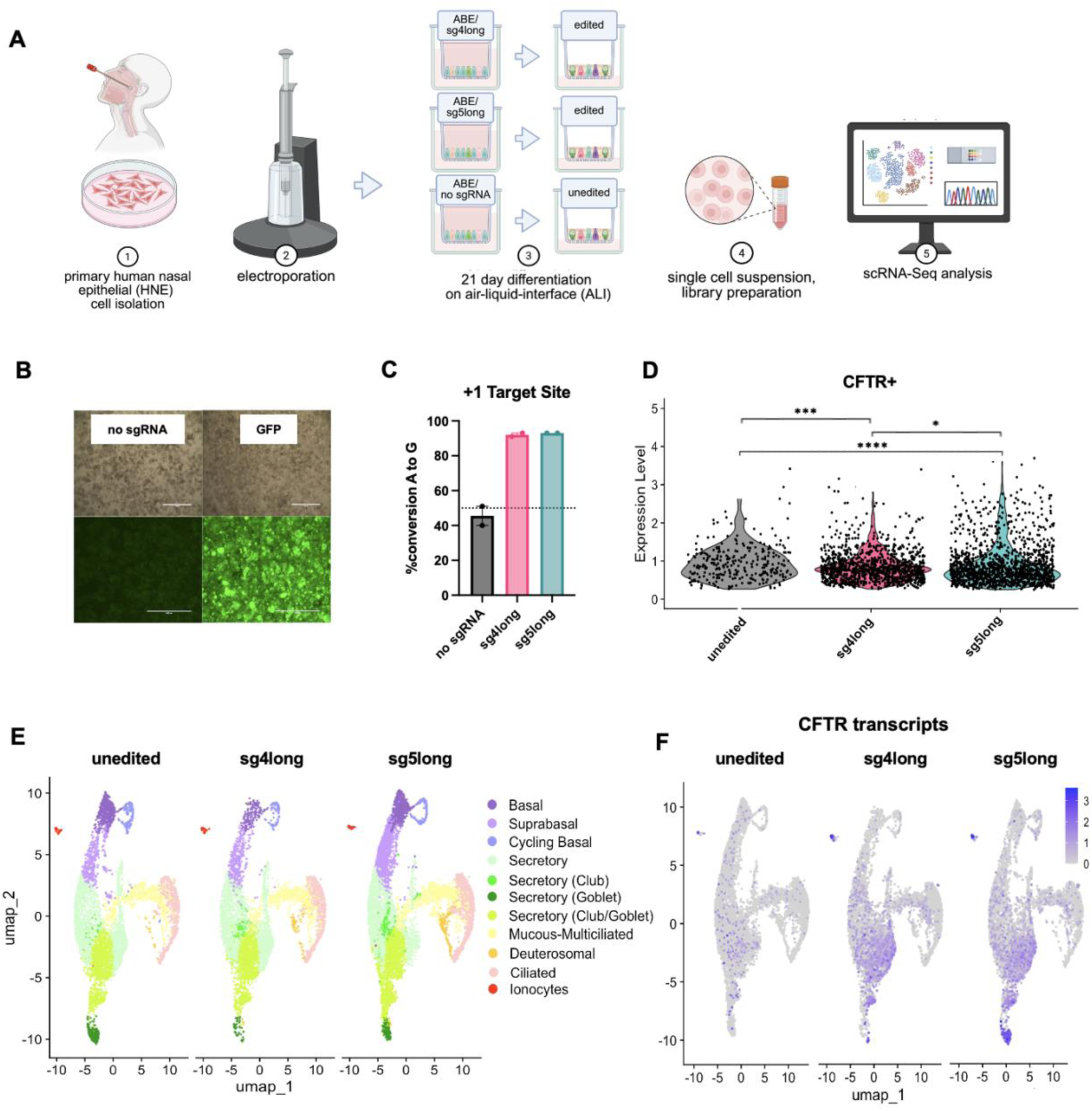
Single-cell transcriptome atlas of primary nasal epithelial (HNE) cells from a CF donor bearing the variants 3120+1G>A/G480S corrected with an adenine base editor. **(A)** Schema of primary HNE collection, electroporation, and differentiation period on air-liquid interface. Cells underwent electroporation and delivery of ABE8e and sgRNA (4long or 5long) before undergoing differentiation for 21 days, then processed for single cell RNA sequencing. **(B)** Fluorescent microscopy images taken 24 hours post electroporation of ABE8e mRNA/no sgRNA (left) and GFP mRNA (right) with brightfield (top) or GFP (bottom). Scale bar is 1000μm. **(C)** Quantification of gDNA A to G nucleotide conversion (%G) at the 3120+1G>A target site. Dashed line representing 50% G nucleotide content to indicate contribution of G sequence from the G480S allele in trans. Values were determined using the Sanger sequencing deconvolution program EditR.[62] Data shown as mean ± SEM (N=2). **(D)** Violin plot comparing CFTR expression levels in CFTR+ cells only across edited and unedited samples. p value was determined by unpaired t-test. ****p ≤ 0.0001. **(E)** UMAPs separated by edited (N=4 (sg4long N=2, sg5long N=2); 13,305 cells) or unedited (N=2 (control); 9,906 cells) samples. **(F)** UMAP detailing CFTR expression across all cellular subtypes.

An integrated uniform manifold approximation and projection (UMAP) assembled using all 23,211 cells revealed 11 distinct cellular subsets including basal, secretory club and goblet, ciliated and ionocytes (**Figure S1**) using marker genes for each subset characterized in prior publications (**Figure S2, S3)**.[12,18,20,31] Unedited and edited CF cells had higher proportions of secretory and ciliated cells but lower fractions of basal cell types than nasal cultures derived from healthy (wild type; WT) individuals (**Figure S4A**), as reported by others.[17,20] All major subclusters were present in projections of unedited (9,906 cells), edited with sg4long (6,231 cells) and edited with sg5long (7,074 cells) (**Figure 1E**). At this level of resolution, ionocytes were differentiated from tuft/brush cells (**Figure S5**), the proposed progenitor of ionocytes.[32,33] The UMAPs revealed that editing with either guide RNA resulted in similar substantial increases in number and similar distribution of cells with detectable *CFTR* transcript (**Figure 1F**). Transcription of the *CFTR* gene bearing G480S accounted for the *CFTR* mRNA in the unedited cells.

### Editing increases detectable CFTR expression in ionocytes and secretory cells resulting in CFTR function increases to clinically relevant levels

To assess editing efficiency in cells endogenously expressing *CFTR* (*CFTR*+), we probed individual cellular subsets. Most subsets were found to have similar proportions between edited and unedited samples, excluding the basal and secretory populations where both editing with either guide RNA resulted in a lower percentage of basal and higher percentage of secretory subsets (**Figure 2A and S6A**). Both edited samples revealed a different *CFTR*-expressing cellular makeup from unedited cells. Notably among all *CFTR*+ cells, the fraction attributable to ionocytes increased by approximately 20% in edited samples (**Figure 2B and S6B**) due to a nearly 5-fold increase in ionocytes with detectable *CFTR* mRNA (**Figure 2C and S6C**). Notably, expression level per ionocyte did not increase in the cells edited with either guide RNA (**Figure 1D**). However, the greater number of ionocytes with detectable levels of *CFTR* resulted in an increase in total % *CFTR* expression attributable to ionocytes with each guide RNA (**Figure 2E**) that translated to a ∼26% increase (from 4.7% to 5.8%) when results from each guide RNA were combined (**Figure S6D**).

**Figure 2:**
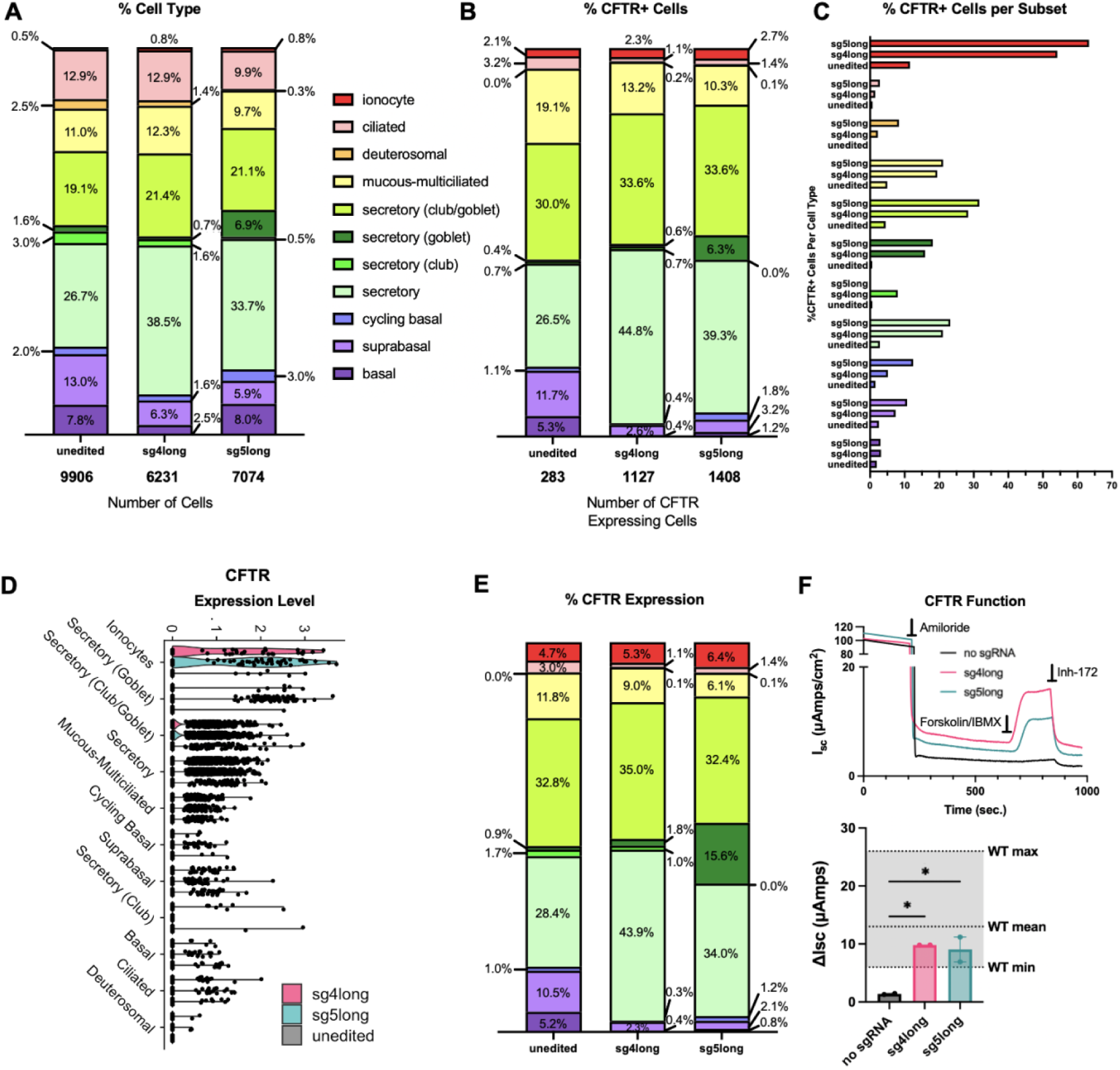
Graphical representations of cell type distribution and CFTR upregulation, CFTR distribution and expression between primary HNE samples unedited (no sgRNA) or edited with sg4long or sg5long. **(A)** Cellular composition across cellular subsets. **(B)** Percent CFTR+ cells across cellular subsets. **(C)** Percent CFTR+ cells per cellular subsets. **(D)** Violin plot comparing CFTR expression levels across cellular subsets. **(E)** CFTR expression across cellular subsets, CFTR+ cells only. **(F) Top:** Representative short circuit current tracings of CFTR functional recovering in primary HNEs bearing the 3120+1G>A/G480S variants after electroporation of ABE/no sgRNA (unedited; black), ABE/sg4long (pink), ABE-sg5long. Aligned according to same baseline after addition of Amiloride. **Bottom:** Quantified CFTR functional recovery as delta Isc (ΔIsc, μAmps/cm^2^). Grey shading indicates ranges of values observed in WT/WT HNE cells as previously reported.[35] Data shown as mean ± SEM (two biological replicates from one transfection). p values were determined by one-way ANOVA followed by Dunnett’s multiple comparisons test. *p ≤ 0.05.

Generally, editing was associated with an increase in secretory populations and decrease in progenitor and ciliated. However, there were exceptions in subtypes; cycling basal cells had a 0.3% and 0.1% increase in cell type and *CFTR*+ cell composition, respectively. Inversely, secretory club cells had a 2.8% and 0.4% decrease in cell type and *CFTR*+ cell composition, respectively (**Figure S6A, B**). Most cell types saw dramatic increases in the fraction of cells with detectable *CFTR* (**Figure 2C and S6C**) while *CFTR* expression level in each cell type remained about the same when comparing unedited and edited samples (**Figure 2D**). As expected, the shifts in cell populations resulted in similar shifts in the % *CFTR* expression; secretory cells accounted for a larger fraction while progenitor and ciliated cell types accounted for a reduced portion (**Figure 2E and S6D**). The two sgRNAs exhibited slight differences. Goblet cells, like ionocytes, had high levels of *CFTR* expression per cell. Editing with sg5long increased the number of goblet cells with detectable *CFTR* mRNA resulting in this cell type accounting for ∼16% of *CFTR* expression after editing, but only a modest increase of 0.9% with sg4long (**Figures 2E**). Inversely, the fraction of *CFTR* expression attributable to secretory cells with sg4long increased from ∼28% to ∼44% after editing while sg5long increased to just ∼34% (**Figure 2E and S6D**). Both samples of edited cells had lower fractions of basal cells, constituting a small fraction of *CFTR* + cells after editing. Individual values for percent *CFTR*+ cells from each subset, *CFTR* expression levels, percent cellular makeup, and percent *CFTR*+ cellular makeup is shown in **Figures S7A-D**.

To determine if editing shifted cell types toward a wildtype (i.e., non-CF) distribution, we compared results with previously analyzed scRNA-Seq dataset from healthy HNEs cultured in the same manner and at the same timepoint of differentiation. Secretory cells accounted for larger fractions of cells with detectable *CFTR* expression and overall % *CFTR* expression in the CF samples (**Figure S4B and C**), which aligns with previous reports.[17,20] Due to the method of data analysis in the WT samples, we could not define as many secretory cell subsets as our prior analysis. However, proportions of cells remained directly comparable since both samples underwent processing at 21 days on ALI culture. Notably, ionocytes were increased and basal cell types were decreased in *CFTR*+ populations and in overall %*CFTR* expression in CF samples. Editing increased the ionocytes fraction but further reduced the fraction of basal cells among *CFTR*+ cells and as % of overall *CFTR* expression. Others have reported similar results when full-length WT *CFTR* is expressed as a transgene in CF airway cells.[34]

CFTR functional recovery was assessed by short circuit measurement in Ussing chambers as previously described.[35] After addition of amiloride to inhibit the epithelial Na+ channel (ENaC), CFTR activity was stimulated using a combination of forskolin and IBMX (3-isobutyl-1-methylxanthine) and CFTR chloride transport was measured as the change in current in response to CFTR-specific inhibitor, inh-172 (**Figure 2F; top**). The increase in current (ΔIsc) in primary cells edited with either guide RNA (sg4long: 9.8 ± 0.01 μAmps A/cm^2^; sg5long: 9.1 ± 2.1 μAmps A/cm^2^) was significantly higher than unedited cells (1.4 ± 0.1 μAmps A/cm^2^) and reached ranges previously reported in WT HNE cells [35] (**Figure 2F; bottom**). The substantial increase in current in edited cells is consistent with the increased the number of airway epithelial cells with detectable *CFTR* transcript in most cell types, particularly ionocytes and secretory cells.

### Polymeric NPs delivering ABE/sgRNA to immortalized CFBEs bearing the 3120+1G>A variant achieve clinically significant CFTR functional recovery

Once we had demonstrated that *CFTR* expression was recovered in the cell types essential for CF treatment, we tested the efficiency of editing upon delivery of the base editor RNA using polymeric NPs (PNP), a clinically viable approach as opposed to electroporation. Our first step was to assess delivery and editing efficiency after PNP delivery to immortalized CFBEs bearing the integrated *CFTR* expression mini-gene with 3120+1G>A (**Figure 3A**). PNPs encapsulating NRCH-ABE8e mRNA and the sg4long RNA [2]were tested over a range of RNA concentrations per PNP formulation (5, 7.5 and 10 μg RNA) and PNP volumes (1, 2, 5, 10 and 20 μl) resulting in delivered RNA ranging from 50 to 1000 μg. Sg4long was chosen due to its ability to achieve higher functional recovery of CFTR, likely due to reduced bystander editing.[2] The highest DNA editing efficiency (7%) was observed at the lower doses (500 ng and 1000 ng), while 5% efficiency was noted at the two highest doses (1500 ng and 2000 ng) (**Figure 3B**). Furthermore, the 1000 ng dose had slightly reduced editing at the +3 bystander site (**Figure 3C**), which is known from previous work to cause minimal (∼20%) exon 18 skipping, resulting in the same aberrant splice isoform generated by 3120+1G>A.[2] CFTR function assessed by short circuit current after forskolin activation and inh-172 inhibition (**Figure 3D; left**) revealed unedited cells showed minimal current (0.9 ± 0.1 μAmps A/cm^2^), whereas PNP transfected cells with ABE8e mRNA and sgRNA generated a response between 23 ± 5 and 39 ± 2 μAmps/cm^2^, all within the 10-25% WT therapeutic range (**Figure 3D; right**).

**Figure 3:**
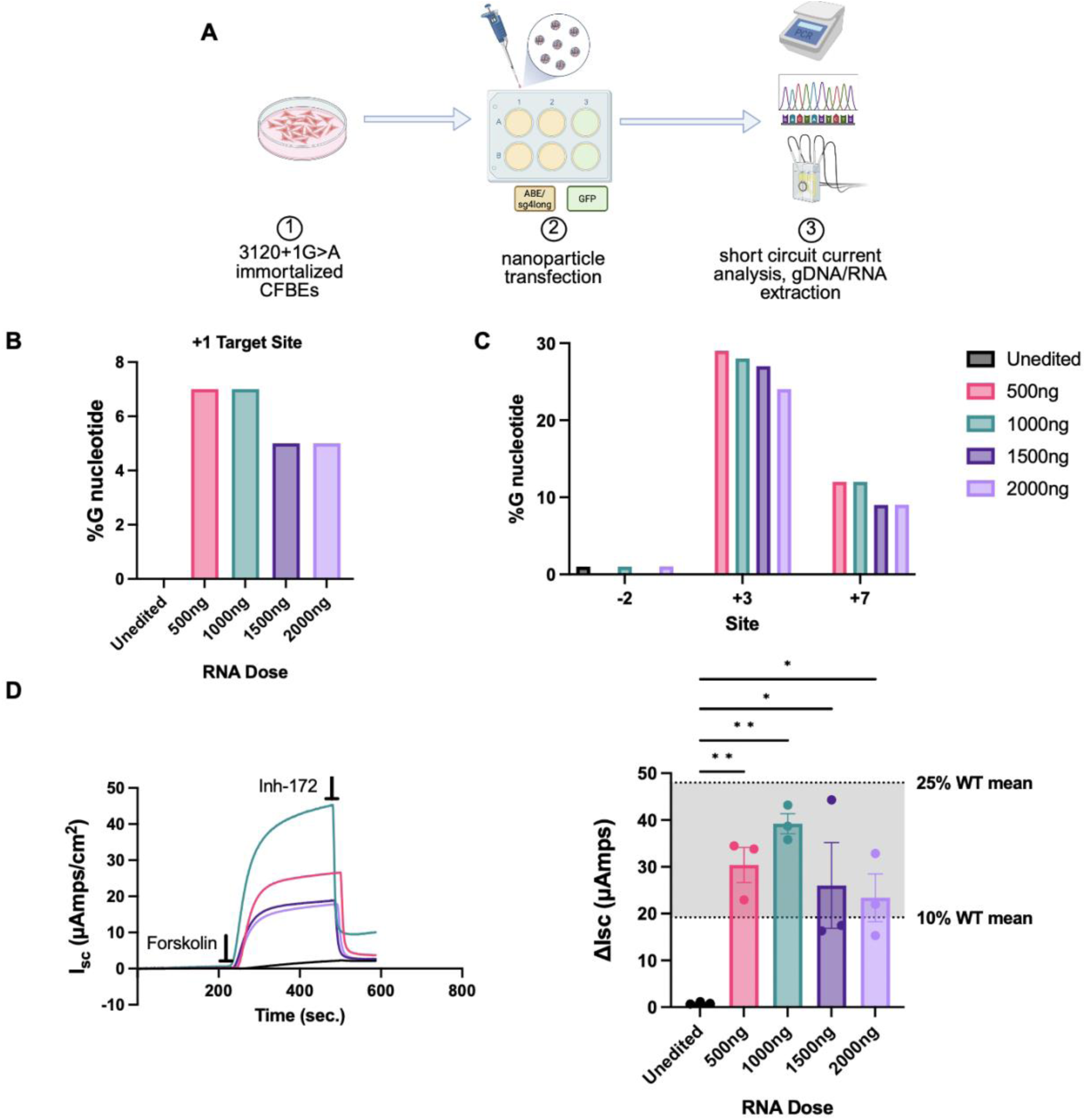
PBAE-E63 nanoparticles delivering ABE8e/sg4long correct 3120+1G>A variant and restore CFTR function in isogenic CFBE cells. **(A)** Schema of immortalized CFBEs bearing the 3120+1G>A variant transfected with PBAE-E63 encapsulating either ABE8e/sgRNA4long or GFP mRNA before analysis by short circuit current analysis, gDNA and RNA extraction. **(B)** Quantification of gDNA nucleotide conversion at the 3120+1G>A (+1) target site, **(C)** as well as potential bystander adenine edits (-2, +3, +7) after delivery of ABE and sgRNA4long to CFBE cells stably expressing EMG_i14-i18 bearing the 3120+1G>A variant. Values determined by Linear/PCR Sequencing using Oxford Nanopore Technology. **(D) Left:** Representative short circuit current tracings of CFTR functional recovering in isogenic CFBEs bearing the 3120+1G>A variant of interest after delivery of GFP (unedited; black), 500ng (pink), 1000ng (green), 1500ng (dark purple) or 2000ng (light purple) with PBAE-E63 encapsulating ABE8e/sg4long. **Right:** Horizontal bar indicates timing of application of each compound, which was sequential. Quantified CFTR functional recovery as delta Isc (ΔIsc, μAmps/cm^2^) across four RNA doses. Grey shading indicates ranges of values observed in WT CFBE cells stably expressing EMG_i14-i18 as previously reported.[2] Data shown as mean ± SEM (three biological replicates from one transfection). p values were determined by one-way ANOVA followed by Dunnett’s multiple comparisons test. **p ≤ 0.01, *p ≤ 0.05.

The 1000 ng dose led to the highest recovery of function, equating to 20% of the function observed in WT CFBE cells, despite the same level of editing at the target site as 500 ng. Earlier studies with GFP mRNA delivery reveal a consistent 80% transfection across all doses while maintaining a viability of >90%, however MFI does decrease as dose increases (**Figure S8**). This indicates there are other factors at play when delivering base editor machinery. Observations with CFBEs noted that delivery with the PNPs was able to correct 3120+1 to clinically significant ranges with relatively low RNA dosing and low rates of editing.

### PBAE PNPs delivering ABE/sgRNA to primary HBEs bearing the 3120+1G>A variant achieve up to 50% WT CFTR functional recovery

Our next step was to assess editing efficiency of PNPs delivered to primary HBEs bearing the 3120+1G>A/F508del variants. Nanoparticles encapsulating NRCH-ABE8e mRNA and sg4longRNA were formulated at four different doses (500 ng,1000 ng, 1500 ng, 2000 ng) and delivered to cells adhered to a 6-well dish. 48 hours post-transfection, cells were moved to filters, where they were allowed to form a confluent monolayer of cells (∼2-3 days), then transitioned to ALI culture to allow for differentiation (**Figure 4A**). Delivery of ABE8e and sgRNA achieved 7.4% correction at 500 ng, 11.3% correction at 1000 ng, 13.4% correction at 1500 ng, and 13.5% correction at 2000 ng doses (**Figure 4B**). After subtracting the contribution from the F508del allele in trans [5] (dashed line, **Figure 4B**), allelic editing was calculated at the 4 RNA concentrations was 15.1%, 22.9%, 27.3% and 27.3%. Bystander editing was noted at the -2, +1, +3, and +7 as previously described,[2] along with editing at the +8 and +9 sites (**Figure 4C**).

**Figure 4:**
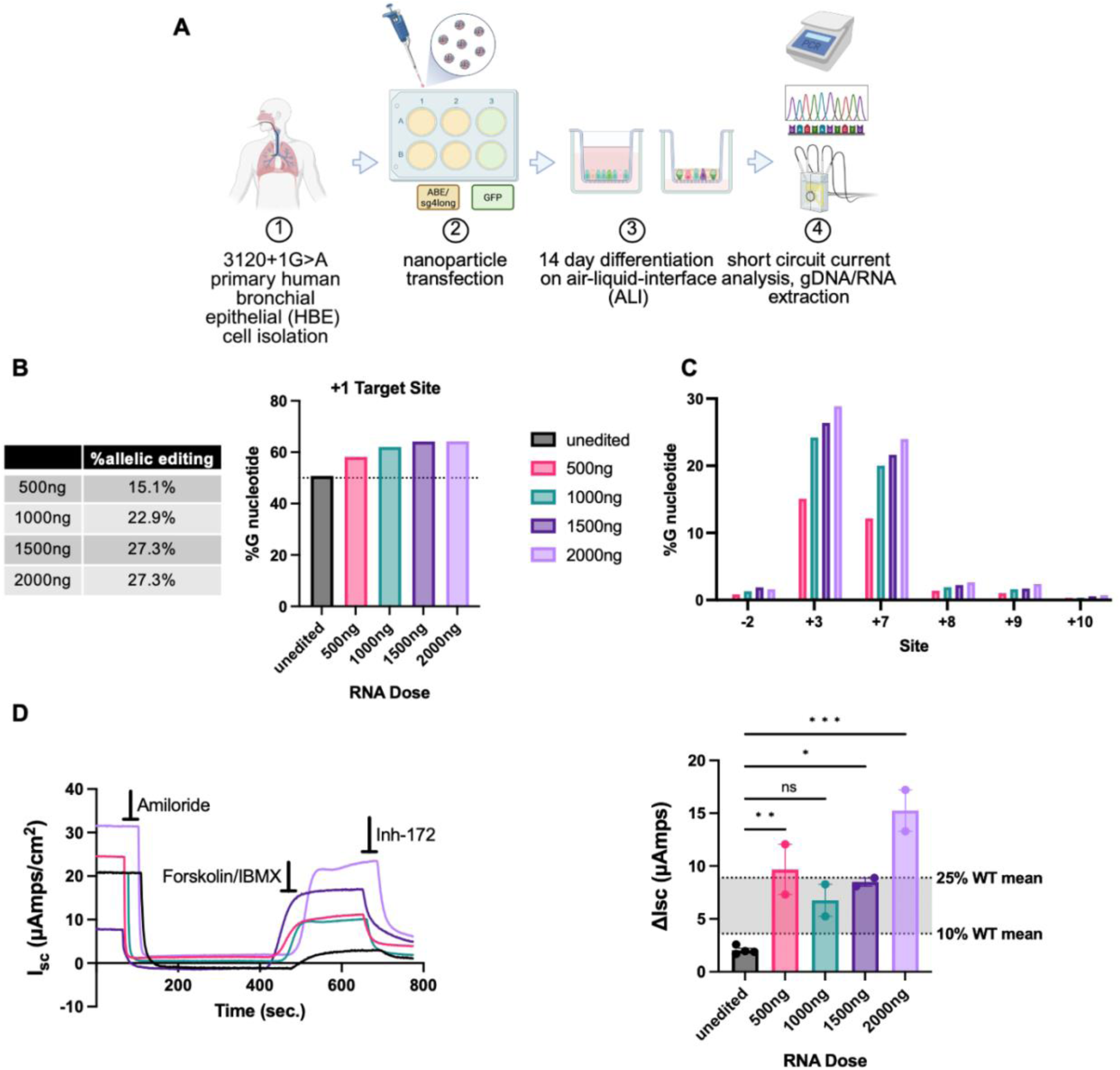
PBAE-E63 nanoparticles delivering ABE8e/sg4long to primary HBE cells bearing the 3120+1G>A/F508Del variants correct 3120+1G>A and restore CFTR function. **(A)** Schema of primary HBE isolation, nanoparticle transfection, and differentiation period on air-liquid interface. Cells were transfected with PBAE-E63 encapsulating either ABE8e/sgRNA4long or GFP mRNA before undergoing differentiation for 14 days, then analysis by short circuit current analysis, gDNA and RNA extraction. **(B) Left:** Levels of allelic editing efficiency under each condition, calculated as: %allelic editing = ((%Gedited-%Gunedited)/%Aunedited)*100.[5] **Right:** Quantification of gDNA nucleotide conversion (%G) at the 3120+1G>A target site (+1). Dashed line representing 50% G nucleotide content to indicate contribution of G sequence from the F508Del allele in trans. **(C)** A to G conversion rate at bystander adenine sites (-2, +3, +7, +8, +9, +10, +11). Values determined using MiSeq and CRISPResso.[64] **(D) Left:** Representative short circuit current tracings of CFTR functional recovering in primary HBE cells bearing the 3120+1G>A/F508Del variants after delivery of GFP (unedited; black), 500ng (pink), 1000ng (green), 1500ng (dark purple) or 2000ng (light purple) with PBAE-E63 encapsulating ABE8e/sg4long. Horizontal bar indicates timing of application of each compound, which was sequential. **Right:** Quantified CFTR functional recovery as delta Isc (ΔIsc, μAmps/cm^2^) across four RNA doses. Grey shading indicates ranges of values observed in WT/WT HBE cells as previously reported.[2] Data shown as mean ± SEM (two biological replicates from one transfection). p values were determined by one-way ANOVA followed by Dunnett’s multiple comparisons test. ***p ≤ 0.001, **p ≤ 0.01, *p ≤ 0.05, p > 0.05.

CFTR function was measured by short circuit current on fully differentiated HBE cells (14 days on ALI culture). As noted above, CFTR chloride transport was defined as the change in current in response to CFTR-specific inhibitor, inh-172 (ΔIsc) (**Figure 4D; left**). Unedited cells showed minimal current (2.0 ± 0.2 μAmps /cm^2^), whereas cells transfected with PNPs ABE8e mRNA and sgRNA generated a response between 7 ± 2 and 15 ± 2 μAmps /cm^2^, with the highest dose equating to 50% of the function observed in WT/WT primary HBE cells (**Figure 4D; right**).[2] This level of function is directly comparable to results achieved by electroporation of the same cargoes in previous studies,[2] and is consistent with our observation above that editing substantially increases the number of *CFTR* expressing ionocytes and secretory cells, the primary mediators of chloride transport.

### Vitronectin is the most abundant protein in the protein corona of PBAE-E63 PNP

We have previously shown that PNP PBAE-E63 can transfect airway epithelial cells after systemic administration to mice.[29] Previous literature has suggested that the protein coating of NPs can enhance delivery to specific organs following systemic delivery.[4] To investigate the constituents of protein coronas, we performed mass spectrometry on PNPs incubated in mouse blood serum. Vitronectin was identified as the protein of highest abundance in PBAE-E63 (**Figure 5A**), as well as cardiac-targeting PBAE-456 (**Figure 5B**). In comparison, non-cardiothoracic targeting PBAE-78018 designed for immune cell transfection does not contain vitronectin among the top 20 proteins (**Figure 5C**). While vitronectin was detected as the most abundant protein for PBAE-E63 (30.6%), levels did not differ substantially from other proteins detected in the coronas. This result is in contrast to Sun et al,[4] where vitronectin was uniquely abundant (∼50%) in the corona of lipid NPs (LNP). Additionally, the top 20 protein composition of lung-targeting PBAE-E63 PNPs differed from that of lung-targeting SORT-LNPs. The PBAEs had significantly more apolipoproteins present and fewer coagulation and immunoglobulin proteins. The difference in overall top 20 protein composition between PBAE PNPs and LNPs also emphasizes the importance of overall protein corona composition as an important biomarker for potential organ and cell-specific targeting.

**Figure 5:**
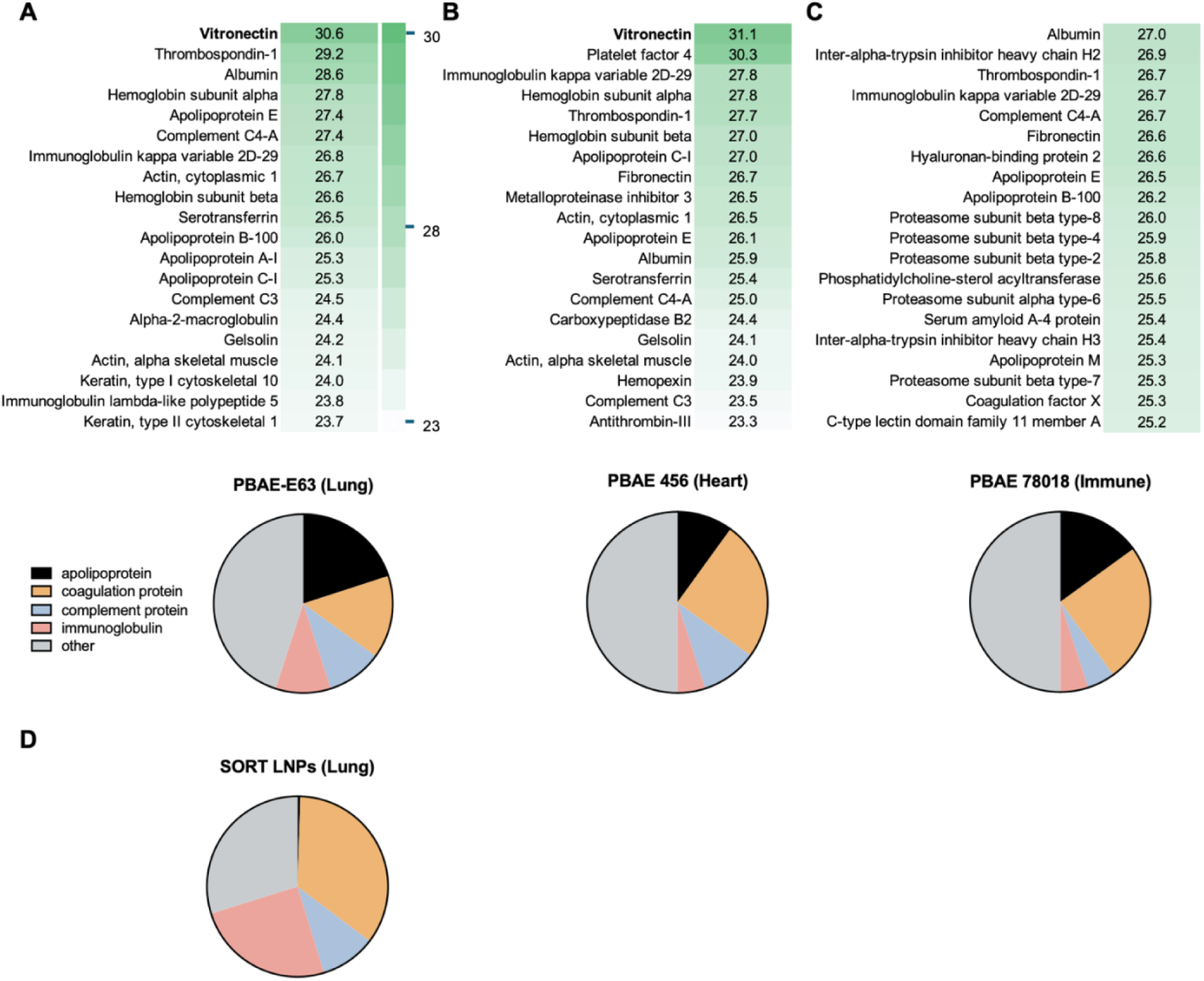
Mass spectrometry proteomics data detailing PBAE NP protein corona compositions. **Top:** Heat map depicting the top 20 most abundant proteins in **(A)** lung targeting PBAE-E63, **(B)** heart targeting PBAE-456, **(C)** immune cell targeting PBAE-78018. **Bottom:** Top 20 proteins classified into physiological categories. **(D)** Top 20 proteins for lung SORT LNP protein corona from *Figure S14A*, Sun Y. et al., Science 2024 [4] classified into physiological categories.

### PNPs can transfect fully differentiated primary HBE 3120+1G>A/F508Del cells including basal cell populations

We next tested whether PNPs transfect fully differentiated human primary airway cells in culture and whether vitronectin coating could enhance the process. PNPs encapsulating GFP mRNA coated at various ratios with either spleen-targeting b2-glycoprotein I (B2) or lung-targeting vitronectin (Vtn) were used to transfect immortalized and differentiated primary HBEs. The highest protein-to-polymer ratio was based on previous literature suggestions that this improved uptake.[4] Flow cytometry revealed that transfection rates of immortalized CFBE cells were similar (80-85%) when different doses (50 ng, 100 ng) of uncoated or B2-coated (low and high) PNPs were used (**Figure 6A**). Uncoated transfection levels were comparable to previously published results with this PNP formulation.[29] In contrast, vitronectin-coated PNPs had an overall decrease in transfection, attaining a maximum of 65%.

**Figure 6:**
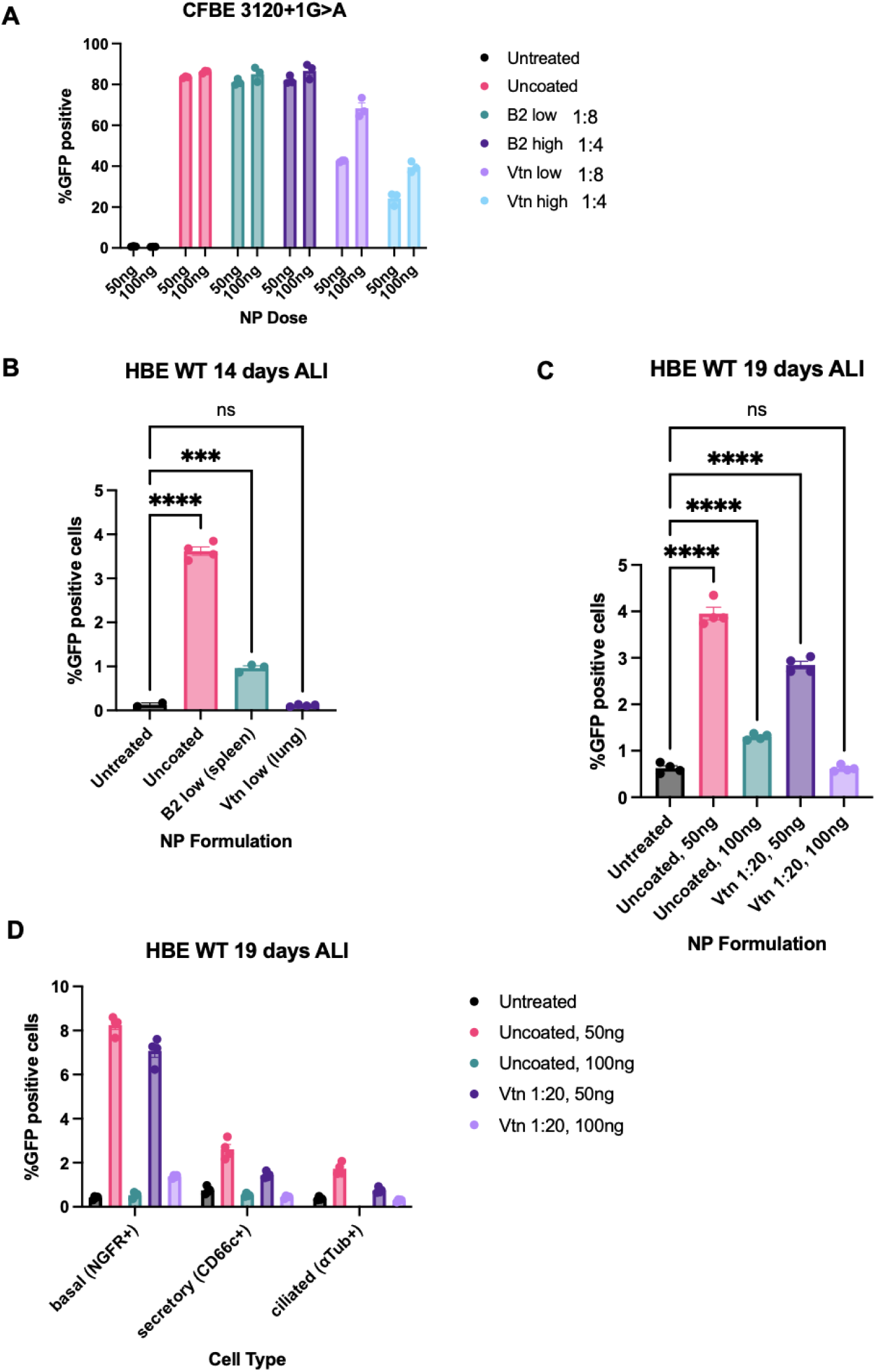
Protein-coated nanoparticle transfection with PBAE-E63, b2-glycoprotein I (B2), and vitronectin (Vtn). **(A)** Flow cytometry results of %GFP positive immortalized CFBE cells bearing the 3120+1G>A variant at two doses (50ng, 100ng) across four treatment conditions (untreated, uncoated, B2-coated, vtn-coated) and two protein-to-PBAE coating ratios (low = 1:8, high = 1:4). Data shown as mean ± SEM (N=3). **(B)** Visualized %GFP positive results from flow cytometry on 14 day differentiated HBE WT cells. Data shown as mean ± SEM (N≥2). **(C)** Percent GFP positive cells from flow cytometry on 19 day differentiated HBE WT cells at 1:20 protein-to-PBAE ratio. **(D)** Percent GFP positive cells in lung cell populations (basal, secretory, ciliated) across the four NP formulation groups. Data shown as mean ± SEM (N=4). p values were determined by one-way ANOVA followed by Dunnett’s multiple comparisons test. ****p ≤ 0.0001, ***p ≤ 0.001, **p ≤ 0.01, *p ≤ 0.05, p > 0.05.

Differentiated primary cells remain difficult to transfect. Given that the lower protein-to-polymer ratio demonstrated better transfection rates in immortalized airway cells, uncoated and coated PNPs were added to primary WT HBEs differentiated on ALI for 14 days. Flow cytometry revealed that uncoated PNPs demonstrated the highest transfection rate of 3.6% in differentiated primary HBEs, with B2-coated NPs demonstrating 1.0%, and vitronectin-coated NPs exhibited no transfection (**Figure 6B**). Due to DLS evidence that vitronectin coated NPs were aggregating at 1:4 and 1:8 ratios, the protein-to-polymer ratio was lowered to 1:20 and NPs were delivered to primary HBE WT cells differentiated for 19 days on ALI. Cells were antibody-stained to elucidate the three major lung cell populations (ciliated, secretory, basal; **Table S1**) and GFP expression was measured by flow cytometry (**Figure S9 and S10**). Uncoated and vitronectin-coated PBAE-E63 PNPs at the 50ng dose had overall transfection rates of 4% and 3%, respectively, whereas higher doses of NPs were less effective (**Figure 6C**). A similar pattern was seen when sorted by cell types. Of note, basal cells achieved the highest rates of transfection of 8% and 7% respectively, with secretory and ciliated cells having lower levels of transfection (**Figure 6D**). These results indicate that differentiated airway cells including basal cells can be transfected with PNPs, but rates are sensitive to dose and vitronectin to polymer ratio.

## Discussion

Gene editing provides an opportunity to cure genetic diseases by correcting deleterious DNA variants *in situ*. In this study, we have addressed several gaps in transitioning gene editing for cystic fibrosis to clinical trials; are cell types essential to pathophysiology corrected, does correction affect cell type distribution and can synthetic vectors such as nanoparticles deliver cargo to essential cell types? As these are pre-clinical studies, we utilized well characterized primary and immortalized human cells in culture to provide a reasonable approximation of differentiated epithelia lining the small airways,[18] a major site of disease in CF.[36–38] scRNA-seq revealed successful correction across a broad range of epithelial cells, including those intimately involved in CFTR-dependent ion transport. We show that progenitor basal cells were edited and successfully transfected in non-differentiated and differentiated states using PNPs.

To our knowledge, this study provides the first single-cell transcriptomic analysis of primary human airway cells after base editing to correct *CFTR*. Correction of the 3120+1G>A variant led to a striking 4- to 7-fold increase in cells with detectable *CFTR* expression, far exceeding the expected two-fold maximum increase due to the monoallelic repair. This disproportionate upregulation occurred across all differentiated epithelial cell types, particularly ionocytes and secretory cells, that are crucial for CFTR-mediated ion transport. CFTR chloride transport increased 6- to 7-fold in edited cells reaching clinically relevant ranges. This functional improvement closely correlated with the increase in overall *CFTR* mRNA expression, consistent with our previous findings that *CFTR* mRNA levels and CFTR chloride transport are linearly correlated in isogenic cell lines.[39] Notably, CFTR function exceeded 10% of WT levels, a threshold generally recognized as sufficient to ameliorate life-limiting lung disease.[40–42]

The increase in *CFTR* expressing cells was consistent across two biologic replicates using two independent guide RNA designs. Small differences in cellular patterns between the sgRNA designs can likely be attributed to a technological rather than biological difference (**Figure S7**). A technical limitation of scRNA-seq may also account for the increase in detectable *CFTR*-expressing cells.[43,44] Indeed, cell types with the highest expression per cell of *CFTR* in unedited samples, such as ionocytes and goblet cells, showed the greatest increase in the number of cells with detectable *CFTR* transcript after editing. There are also several possible biological explanations for this disproportionate increase. One is that the promoter and/or enhancer elements driving expression of *CFTR* bearing the 3120+1G>A allele are considerably more potent than those in the accompanying gene bearing G480S. However, the average *CFTR* expression level per *CFTR*+ cell was not higher in the edited compared to unedited cells. Another explanation is that allelic correction increased the growth rate of *CFTR*-expressing cells, which paralleled the overall increase in *CFTR*+ cells. However, we did not observe substantial differences in growth rates between edited and unedited cells. [43,44] In sum, our results indicate that gene correction may have a disproportionately large impact on overall *CFTR* expression, potentially enhancing functional recovery beyond what would be predicted for a single allelic correction, though the mechanism of this observation remains unknown.

We expected correction of *CFTR* would shift the developmental pattern and cellular composition of differentiated cultures, possibly to that observed in WT individuals. Following editing, we observed a modest change in the relative proportions of major cell types, with an increase in secretory cells and decrease in basal cells. Notably, overall nasal cell distributions in edited and unedited CF cells were more similar to each other than nasal cells derived from healthy individuals. These findings align with two recent studies that reported modest changes in bronchial epithelial cell distributions after transduction of basal cells with lentivirus [34] or AAVs integrating *CFTR* cDNA.[45] Although nasal epithelial cell culture models differ from freshly acquired tissues,[22,46] they still effectively recapitulate the primary role of secretory cells in overall *CFTR* expression,[18] as well as the high per-cell *CFTR* expression in ionocytes and goblet cells.[11,12] Thus, while recovery of *CFTR* function in airway cells from individuals with CF does not recapitulate a wild-type pattern, it does not drastically distort the essential cellular populations needed for a functional differentiated epithelium.

Long-term treatment of genetic disorders requires correction of deleterious variants in progenitor cells. It is widely accepted that basal cells can differentiate into many, or possibly all, respiratory epithelial cell types,[19] a process recapitulated when primary airway cells are cultured at ALI.[47,48] In our study, basal cells, particularly suprabasal and cycling basal cells, were edited as indicated by the increase in *CFTR*+ cells. It has been reported that basal cell transfection does not impact progenitor capacity [34] or drastically alter chromatin structure upon editing.[45,49] However, we observed a consistent reduction in the proportion of basal cells, both overall and among *CFTR*+ cells, after editing. This decrease in basal cell fractions exacerbated differences from healthy ALI culture observed in our study and reported by others.[17,50] Additionally, the only secretory cell subtype to experience a decrease between unedited and edited samples were secretory club, which contain a progenitor capacity similar to basal cells. Whether this shift in progenitor cell populations could affect long-term reconstitution of the airway epithelium remains an important question that could be addressed by longitudinal study of airway epithelial changes in individuals treated with CFTR modulators. Additionally, future studies to track cell fate could determine whether the reduction of basal cells is due to enhanced differentiation into secretory cell subtypes. The most significant increase in *CFTR* within a cellular subset were ionocytes, with the largest overall increase being in secretory cell populations, potentially informing the most ideal targets for CFTR correction and functional recovery.

Given the promising functional response and evidence of widespread editing across key epithelial cell types, we explored the potential of a clinically viable delivery method. Our group had previously shown that polymeric NPs (PNP) efficiently deliver RNA encoding GFP and Cre mRNA reporters to airway epithelial cells following systemic injection.[29] Although correction in isogenic immortalized CFBE cells reached modest levels (5-7% at the target nucleotide), CFTR function exceeded the 10% WT threshold. Interestingly, target base editing after PNP transfection was considerably higher in the primary cells (15-27%) compared to immortalized isogenic cells. Despite lower editing frequencies, PNP-mediated delivery of the editing machinery achieved similar levels of CFTR function as electroporation. One possible explanation is that the PNPs may be less toxic than electroporation, as previously reported.[51] In addition, our *in vitro* results suggest that even low levels of CFTR correction may be sufficient for reversal of the CF phenotype.[52]

Despite encouraging evidence that modest correction may result in clinically relevant outcomes, it remains desirable to maximize the uptake of delivery vehicles such as NPs. In the case of lipid NPs (LNP), vitronectin has been proposed as a key protein for *in vivo* targeting of lung SORT LNPs.[4,53,54] Thus, it was notable that vitronectin was the most abundant component of the lung-targeting PNP protein corona used here. However, vitronectin and albumin are the only proteins in the top 20 most abundant that both lung-targeting NP classes have in common, with physiological characteristics of the other abundant proteins varying greatly, suggesting that a targeting mechanism might lie in multiple contributing factors.[55,56] Notably, vitronectin was also the most abundant protein present on the protein corona of cardiac-targeting PNPs. Coating of PNPs encapsulating GFP mRNA with vitronectin did not improve transfection efficiency of either immortalized CFBE cells or fully differentiated primary airway cells. Our result is in contrast with that reported for lung targeted SORT-LNPs, which may be due to the much higher abundance of vitronectin (∼50% of protein) on lung targeted SORT-LNPs, possibly suggesting a greater avidity of the LNP corona for vitronectin due to a more neutral surface charge (**Table S2**).[4,57] Despite the lack of vitronectin-enhanced uptake, up to 8% of basal cells were GFP+. Since differentiated airway cultures may be a reasonable proxy for airways *in vivo*, future studies should explore whether other PNP formulations with or without specific corona proteins substantially increase update in key cell types for CFTR correction.

The study has several limitations. A technical constraint of scRNA-seq is the limit of transcript detection. Modest increases in expression may translate into a disproportionate number of cells crossing the detection, thus accounting for a 4- to 7-fold increase in *CFTR* expression.[43,44] We report the reproducibility of this finding and propose explanations at the regulatory and cellular level, but we do not know if this phenomenon is a consequence of correction of a splice site variant. Correction of other variants such as missense variants may have different effects.[11,12,17,18,46] Differences in culture conditions, including days on ALI and media choice both influence cell type and lineage,[22,46,58] and subject to subject variation.[17] The matched comparison used here reduced variability due to these effects. Studies with PNPs highlighted further improvement for both delivery of various cargoes and enhancement of base editor efficiency. Despite scaling the amount of RNA delivered accordingly, allelic conversion rates never exceeded more than 27%, and although this level corrected *CFTR* function to clinical relevance in cell culture, it may be insufficient *in vivo.* Strategies to improve delivery could include optimizing PNP surface chemistry and enhancing editing efficiency with a more recently evolved and thus more accurate base editor to minimize potentially deleterious bystander effects. Finally, it remains unknown how well human primary cell culture translate to *in vivo* efficacy and why corrected primary cell subtype distributions migrated further from that seen in WT cultures.

In summary, these studies demonstrate that modest levels of editing may increase the level of *CFTR* in a disproportionate number of airway cells thereby driving substantial improvement in CFTR-mediated transport to clinical relevance CF. These studies provide key information about the cellular populations that undergo changes in expression when *CFTR* is successful edited as well *in vitro* evidence that inform translatability of PNP-mediated base editor gene therapy to potentially cure CF lung disease.

## Materials and Methods

### Polymer synthesis

Polymers were synthesized using previously reported protocols.[59] Briefly, monomers Bisphenol A glycerolate diacrylate (B7) and Trimethylolpropane triacrylate (B8) were dissolved at 600 mg/mL in dimethylformamide reacted with side-chain monomers (4-amino-1-butanol (S4), 4-(2-aminoethyl)morpholine (S90), and 1-dodecylamine (Sc12)) with 80:20 mol/mol c12:90 with stirring for 48 h at 85°C to allow polymerization via stepwise Michael addition reactions. Monomers were reacted at an overall vinyl:amine ratio of 2.3 to allow acrylate-terminated polymers to form. Polymers were end-capped by further reaction with primary amine-containing E monomers (diethylenetriamine (E63)) at room temperature for 2 hrs [200 mg/mL polymer and 0.3 M E monomer in tetrahydrofuran (THF)] and purified by two diethyl ether washes. Diethyl ether was decanted and the sample was dried thoroughly under vacuum. The synthesized polymer, 7-90,C12-63 (PBAE-E63), was dissolved in dimethyl sulfoxide at 100 mg/mL and stored at -20°C with desiccant in single-use aliquots until used.

### Nanoparticle formulation

For *in vitro* studies, NPs were prepared by dissolving polymer and mRNA at 1 μg/μL separately in 25 mM sodium acetate (NaAc; pH 5), then the two solutions were mixed at 1:1 volume ratio to allow for self-assembly at room temperature (RT) for 10 min.

### Cell Culture

Primary HNE cells (c.3846G *>*A/c.3846G *>*A; P3) were obtained by nasal brushing performed by Dr. Christian Merlo at Johns Hopkins Hospital under IRB no.00116966. Cells were propagated on a feeder layer of 3T3 mouse fibroblasts irradiated with 30 Gy. To allow for expansion, cells were maintained in the presence of 10 mM reagent Y-27632 2HCl (ROCK inhibitor, Selleckchem). Passage number (P1, P2, P3, P4) reflects the timepoint of growth in a 6-well dish or T25 flask before transitioning to filters. Primary HNE cells were grown on a feeder layer of irradiated 3T3 mouse fibroblast cells before being isolated for transfection studies. Human bronchial epithelial (HBE) cells (CF: c.2988+1G *>*A/c.1521_1523delCTT; P3, WT: P3) were obtained through the CFF therapeutics lab biobank. Passage number represents time since being thawed.

Immortalized bronchial epithelial cells from a person with CF (CFBE41o-) [60] with an integrated target for FlpIn recombinase. (CF8Flp cells) [61] into which a *CFTR* expression minigene (EMG) containing 3120+1G>A (c.2988+1G *>*A) had been inserted were used for CFBE cell screens. The minigene contained all exons of *CFTR* and appropriately positioned flanking intron sequences from exons 14 to 18, as previously described.[2] Cells were maintained in Gibco’s minimal essential medium with 10% FBS, 1% PS and 200 mg/mL Hygromycin B to maintain stable integration of the *CFTR* EMG. All cells were maintained at 37°C with 5% CO_2_.

### Transfection

#### Immortalized CFBEs and primary HBE cells

Immortalized and primary HBE CF cells were plated in either 24 or 6-well tissue culture plates and allowed to adhere. Once at ∼80 % confluence, cells were transfected with mRNA-encapsulating NPs. NPs were formulated following the *in vitro* transfection formulation described above; NP solution was added to 2 mL fresh media, then incubated with cells for 2 hrs before undergoing a media change. 48 hours post-HBE transfection, cells were moved to filters for differentiation. Cells remained in medium containing ROCK inhibitor until confluency was reached (∼1–2 days), at which point cells were moved to differentiation medium (no ROCK inhibitor). The next day, apical medium was removed, starting ALI culture. Primary cells were maintained on ALI culture for 14 days before short circuit measurements and collection of gDNA and RNA. Cells transfected with GFP mRNA were collected 24 hours post-transfection and stained with 7-AAD (1:200 dilution) with transfection efficacy evaluated via flow cytometry (Attune NxT flow cytometer, Thermo Fisher Scientific) and analyzed using FlowJo software (FlowJo, Ashland, OR). The expression of GFP was quantified by normalizing the geometric mean fluorescence intensity (MFI) of each NP treatment to MFI of the untreated group. Gating strategies to identify GFP+ cell populations are provided in **Figure S9** and **S10.**

#### Primary HNE cells

Base editor and sgRNA were delivered to non-differentiated primary cells via electroporation. Base editor mRNA was at a concentration of 2 μg/μL and synthetic sgRNA (IDT) was at a concentration of 100 μM. Editor and sgRNA were combined in a 3:1 volume ratio and a total of 1 μL of RNA was used for each electroporation reaction. In the no sgRNA control, synthetic sgRNA was substituted with equivalent volume of DEP-C H_2_O and for GFP control eGFP mRNA with unmodified bases and CleanCap was used (TriLink Biotechnologies, catalog no. L-7601). Electroporation was performed using the Neon Transfection System (Invitrogen) with 10μL Neon tips (Invitrogen, catalog no. MPK1025). Each electroporation reaction consisted of 1.5×10^5^ undifferentiated primary cells resuspended in 9 μL of buffer R (Invitrogen, catalog no. MPK1025) and combined with 1 μL of nucleic acid mix. Electroporation conditions were as previously described.[2] After electroporation, cells were plated onto human collagen type IV (Sigma, no. C6745-1ML) coated snapwell filters (Costar, catalog no. 3801) in a 6-well plate seeded with irradiated mouse fibroblast cells (3T3). Two electroporation reactions were added to each filter. Fluorescence microscopy was performed 24 hrs post-electroporation to validate successful transfection of the GFP control. Cells were allowed to recover from electroporation in medium containing ROCK inhibitor until confluency was reached (∼5–10 days), at which point cells were moved to differentiation medium (no ROCK inhibitor). The next day, apical medium was removed, starting ALI culture. Cells were maintained on ALI culture for 21 days before short circuit measurements and collection of gDNA and RNA.

#### Protein coating

Primary HBE WT cells were grown on a feeder layer of irradiated 3T3 mouse fibroblast cells before being isolated and moved to filters, where they underwent the same timeline of differentiation as described above for 14-19 days before being transfected with NPs. 24 hours post transfection, cells were harvested to create a single cell suspension. Surface staining of cells was then performed with fluorescent antibodies was performed using the antibodies and dilutions listed in **Table S1** in FACS buffer for 30 minutes at 4°C at which time cells were washed twice and resuspended in FACS buffer for analysis using an Attune NxT flow cytometer (Thermo Fisher Scientific). Data were analyzed using FlowJo software (FlowJo, Ashland, OR). Gating strategies to identify cell populations are provided in **Figure S10**. Wells with at least 3,000 of each cell type analyzed and reported from flow cytometry.

### Genomic DNA extraction/PCR amplification/Sequencing

For CFBE cells and primary airway cells, gDNA was collected directly from snapwell filters after short circuit measurements were taken. Genomic DNA (gDNA) extraction was performed using the DNeasy Blood & Tissue Extraction Kit (QIAGEN, catalog no. 69504). Following extraction, editing was assessed by PCR amplification of the relevant *CFTR* exon as previously described.[2] For CFBE experiments PCR was performed with KOD hot start master mix (Millipore Sigma). For primary cells, PCR was performed using HotStarTaq DNA polymerase (QIAGEN, catalog no. 203203). CFBEs underwent Linear/PCR Sequencing performed by Plasmidsaurus using Oxford Nanopore Technology with custom analysis and annotation. Primary HNE PCR products were purified before undergoing Sanger sequenced with editing rates at the target and bystander sites quantified with EditR.[62] Primary HBE editing was assessed via MiSeq (see *High Throughput Sequencing*).

### High Throughput Sequencing

Targeted amplicons were generated using gene-specific primers with partial Illumina adaptor overhangs (Forward primer: 5-ACACTCTTTCCCTACACGACGCTCTTCCGATCTNNNN [gene-specific sequence-3’; Reverse primer: 5’-TGGAGTTCAGACGTGTGCTCTTCCGATCT [gene-specific sequence]-3’) and sequenced as previously described.[63] Gene-specific primer sequences are listed in **Table S3**. Extracted HBE gDNA (as described above) was used as a template to amplify the target site. Amplicons were indexed in a second PCR reaction and pooled for sequencing. Gel purification of the indexing PCR product on a 1% agarose gel was used to purify amplicons from primers. Purified amplicons were quantified using qPCR. 10% PhiX Sequencing Control V3 (Illumina) was added to the pooled amplicon library prior to running the sample on a MiSeq Sequencing System (Illumina) to generate single end 300 bp reads. Samples were demultiplexed using the index sequences, fastq files were generated, and alignments and editing quantification was conducted using CRISPResso2.[64]

### RNA analysis/cDNA synthesis

An input of 500 ng of RNA was used with the iScript cDNA synthesis kit (Bio-Rad). For immortalized cells, cDNA was diluted 1:10.

### Evaluation of CFTR channel function

#### Immortalized CFBE cells

Cells were grown on filters until a transepithelial resistance of >200 Ohms was reached (6–8 days). Filters were then mounted on Ussing chambers, and a chloride gradient was established using asymmetrical buffers as previously described.[2] Short circuit current measurements were taken using a multi-channel voltage-current clamp amplifier (Physiologic Instruments) and the data acquisition program Acquire and Analyze. After equilibration, forskolin (10 mM) was added to the basolateral chamber to activate channel opening. The CFTR-specific inhibitor, inh-172 was used (10 mM, apical chamber) to inhibit the channel. The drop in current after addition of inhibitor allowed for quantification of CFTR function (ΔIsc).

#### Primary cells

CFTR function was assessed in primary cells by mounting differentiated filters on Ussing chambers using a symmetrical buffer (126 mM NaCl, 25 mM NaHCO3, 5 mM KCl, 2.5 mM Na2HPO4, 1.8 mM CaCl2, 1 mM MgSO4, 10 mM dextrose). Buffer pH was maintained at 7.3–7.4 by continuous circulation with carbogen gas (95% O_2_/5% CO_2_) and temperature was maintained at 37°C. The apically located sodium epithelial channel (ENaC) was inactivated by addition of amiloride. A combination of forskolin and IBMX was added to the basolateral chamber to stimulate channel opening. Inh-172 was used to inhibit CFTR channel function and quantify ΔIsc.

### scRNA-seq

Library preparation was performed in the Cutting lab using 10X Genomics v.3.1 30 Library Preparation chemistry with a dual indexing system. Sequencing was performed by the Johns Hopkins GRCF via paired-end sequencing 2×100 cycles using a NovaSeq 6000 Illumina sequencing machine. Analysis of scRNA-seq results was performed using 10X Genomics Cell Ranger 3.1.0 followed by the Seurat package (v.4.2) in R created by the Satija lab.[65] Subsetting was determined based on comparisons of the number of genes expressed, UMI count, and percentage of reads mapped to mitochondrial RNA. A log normalization with a scale factor of 10,000 was used. Cell type was assigned based on published transcriptional markers for each cell type: basal (6.4%; KRT5+, TP63+), cycling basal (2.2%; KRT5+, TOP2A+), suprabasal (9.1%; KRT5+, TP63-), secretory (32.0%; SCGB3A1+), secretory (club) (1.9%; SCGB1A1+), secretory (goblet) (3.0%; MUC5B+), secretory (club/goblet) (20.3%; SCGB1A1+, MUC5B+), mucous-multiciliated (10.9%; FOXJ1+, MUC5AC+), deuterosomal (1.5%; DEUP1+, FOXJ1+), ciliated (12.0%; DEUP1-, FOXJ1+)), and ionocytes (0.7%; FOXI1+, *CFTR*+).[11,12,18,20]

### Protein corona data collection and analysis

Nanoparticles formulated for *in vivo* use as described previously[29] were incubated in mouse blood serum (Bio-Rad) at 200 μL/mg for 1 hour, pelleted by centrifuge, then underwent preparation by Proteomics iST kit (PreOmics). Analysis was performed by liquid chromatography-mass spectrometry (LC-MS). Prior to MS analysis, samples were desalted using a 96-well plate filter (Orochem) packed with 1 mg of Oasis HLB C-18 resin (Waters). Tryptic peptides were analyzed on an a Dionex RSLC Ultimate 300 (Thermo Scientific) coupled to an Orbitrap Exploris 480 Mass spectrometer (Thermo Fisher Scientific). Peptides were separated using a 60-minute gradient from 4-30% buffer B (buffer A: 0.1% formic acid, buffer B: 80% acetonitrile + 0.1% formic acid) at a flow rate of 300 nl/minute. Briefly, for DDA-MS, the full MS scan was set to 300-1,200 *m/z* in the Orbitrap with a resolution of 120,000 (at 200 *m/z*) and an AGC target of 5×105. MS/MS was performed in the ion trap using the top speed mode (2 seconds), an AGC target of 1×104, and an HCD collision energy of 35.

### Graphical illustrations

Graphical illustrations were created using BioRender (https://biorender.com/).

### Statistical analysis

GraphPad Prism v.10.1.1 (GraphPad Software), was used to perform all statistical analysis. Data are shown as mean ± SEM. Unless otherwise stated, the absence of statistical significance markings, where a test was stated to have been performed, signifies no statistical significance. Statistical significance is denoted as follows unless otherwise noted: *P *<*0.05, **P *<*0.01, ***P *<*0.001, and ****P *<*0.0001. n. s., not significant.

## Supporting information

scRNASeq DGE

## Data Availability

Values for all data points in graphs are reported in the Supporting Data Values file. scRNA-seq and Mi-Seq data reported in this article are available from the corresponding author upon reasonable request.

## AUTHOR CONTRIBUTION

Conceptualization was contributed by EWK, ATJ, SYT, GAN, JJG, and GRC. Methodology was contributed by EWK, ATJ, ARP, ACE, and SYT. Investigation was contributed by EWK, ATJ, ARP, ACE, AIP, KLS, MJ, SSS, and SYT. Visualization was contributed by EWK, GAN, and NS. Funding acquisition was contributed by EWK, SYT, NS, JJG, and GRC. Project administration was contributed by JJG and GRC. Supervision was contributed by JJG and GRC. Writing of the original draft was contributed by EWK, JJG, and GRC. Review and editing of the manuscript were contributed by EWK, JJG, and GRC.

## ACKNOWLEDGEMENTS

The authors would like to thank the healthy individuals and individuals with CF for their donation of primary HNE cells and Dr. Christian Merlo at Johns Hopkins for obtaining these samples, and Priyanka Bhatt at the CFF Therapeutics Lab for assistance with shipping and methodology for culture of primary HBE cells. The authors would also like to acknowledge Dr. Simone Sidoli at Albert Einstein College of Medicine for providing the resources to run proteomics sample on Mass Spectrometry. This work was supported by the US National Institutes of Health (R01EY031097 and P41EB028239 to J.J.G., R37CA246699 to S.Y.T.), the Cystic Fibrosis Foundation (004319G222 to E.W.K., S.Y.T., G.R.C., J.J.G., CUTTIN20G0 to G.R.C, SHARMA23GO1 to N.S., NEWBY23XX0 to G.A.N.), the Spruance Foundation and Vertex Pharmaceuticals (Vertex Research Innovation Award to N.S.). E.W.K. thanks the XDBio program for support.

## DECLARATION OF INTERESTS

The authors declare the following financial interests/personal relationships which may be considered as potential competing interests: Jordan Green reports financial support was provided by National Institutes of Health. Garry Cutting reports financial support was provided by Cystic Fibrosis Foundation. Stephany Tzeng reports financial support was provided by National Institutes of Health. Jordan Green reports a relationship with Dome Therapeutics that includes: board membership, consulting or advisory, and equity or stocks. Jordan Green reports a relationship with Cove Therapeutics that includes: board membership, consulting or advisory, and equity or stocks. Jordan Green reports a relationship with WyveRNA Therapeutics that includes: board membership and equity or stocks. Jordan Green reports a relationship with VasoRx that includes: board membership and equity or stocks. Garry Cutting reports a relationship with Cystic Fibrosis Foundation that includes consulting or advisory. Gregory Newby reports that he has filed patents on base editing technology. Johns Hopkins University has filed a patent application based on technology discussed in this manuscript with J.J.G., S⋅Y.T., G.R.C., and E.W.K. as co-inventors. The other authors declare that they have no competing financial interests or personal relationships that could have appeared to influence the work reported in this paper.

