## Supplemental Figures 1-10, Supplemental Tables 1-3 for "Base editing and nanoparticle transfection of airway cell types essential for treatment of cystic fibrosis"

**Supplemental Material**

10 Figures
3 Tables

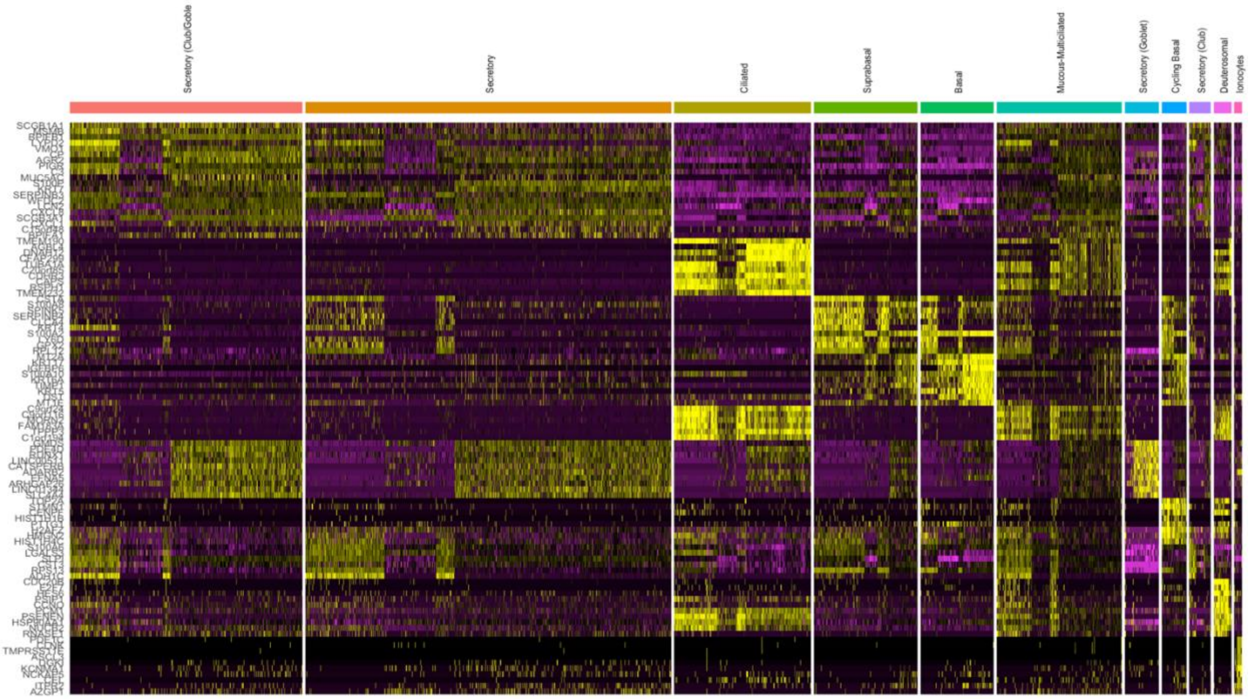


**Figure S1.** Scaled expression of the top DEGs that inform specific cell subsets, for edited and unedited samples further separated by subset, visualized by heat map.

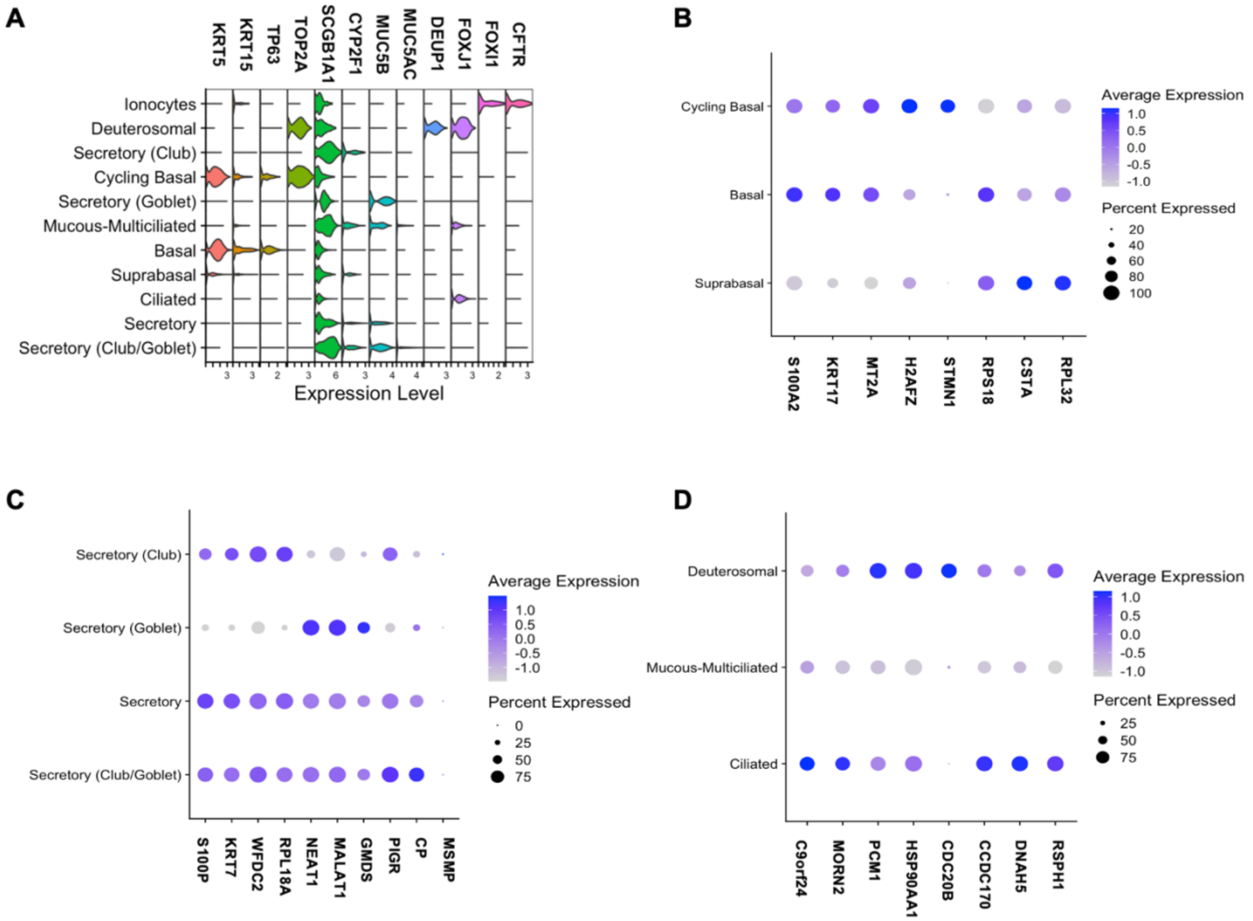


**Figure S2.** Select gene expression across cell subsets. **(A)** Violin plot visualizing top marker genes that inform individual cellular subsets. Comparison of subset-specific gene expression among **(B)** progenitor, **(C)** secretory, and **(D)** ciliated cell populations.

A

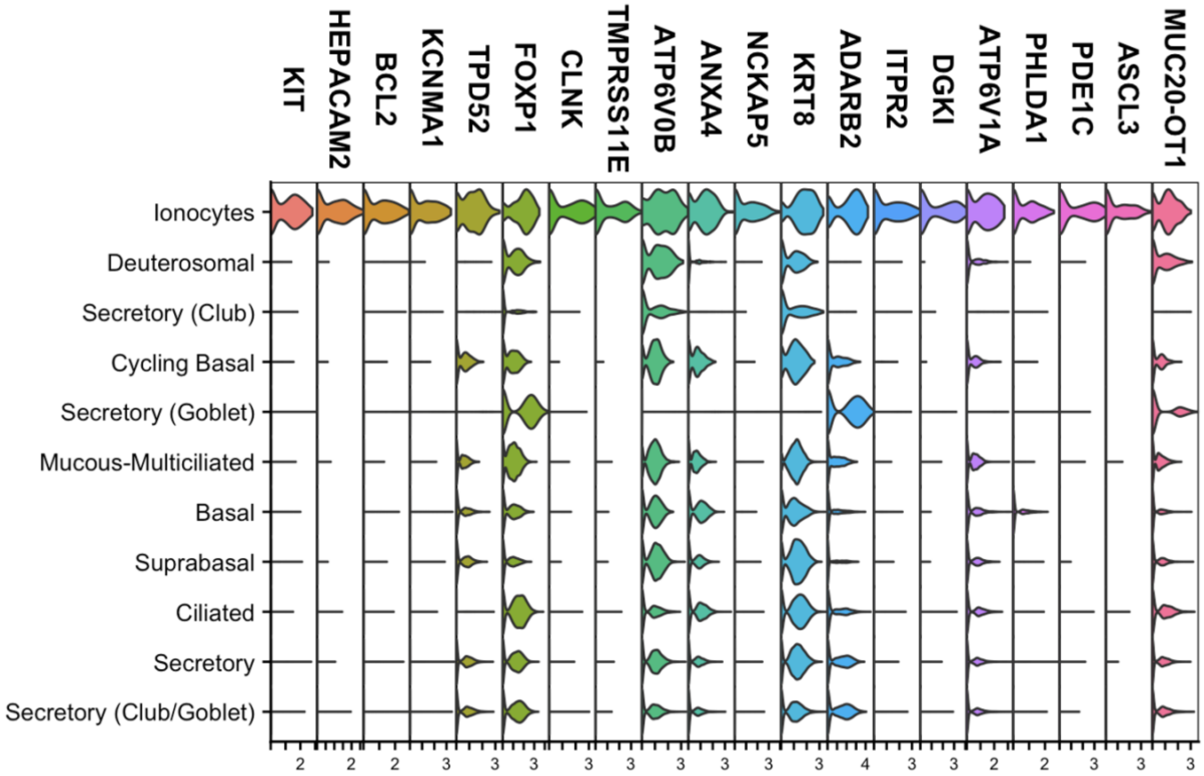

Cluster Markers for Ionocytes
KIT
HEPACAM2
BCL2
KCNMA1
TPD52
FOXP1
CLNK
TMPRSS11E
ATP6V0B
ANXA4
NCKAP5
KRT8
ADARB2
ITPR2
DGKI
ATP6V1A
PHLDA1
PDE1C
ASCL3
MUC20-OT1
Ionocytes
Deuterosomal
Secretory (Club)
Cycling Basal
Secretory (Goblet)
Mucous-Multiciliated
Basal
Suprabasal
Ciliated
Secretory
Secretory (Club/Goblet)

B

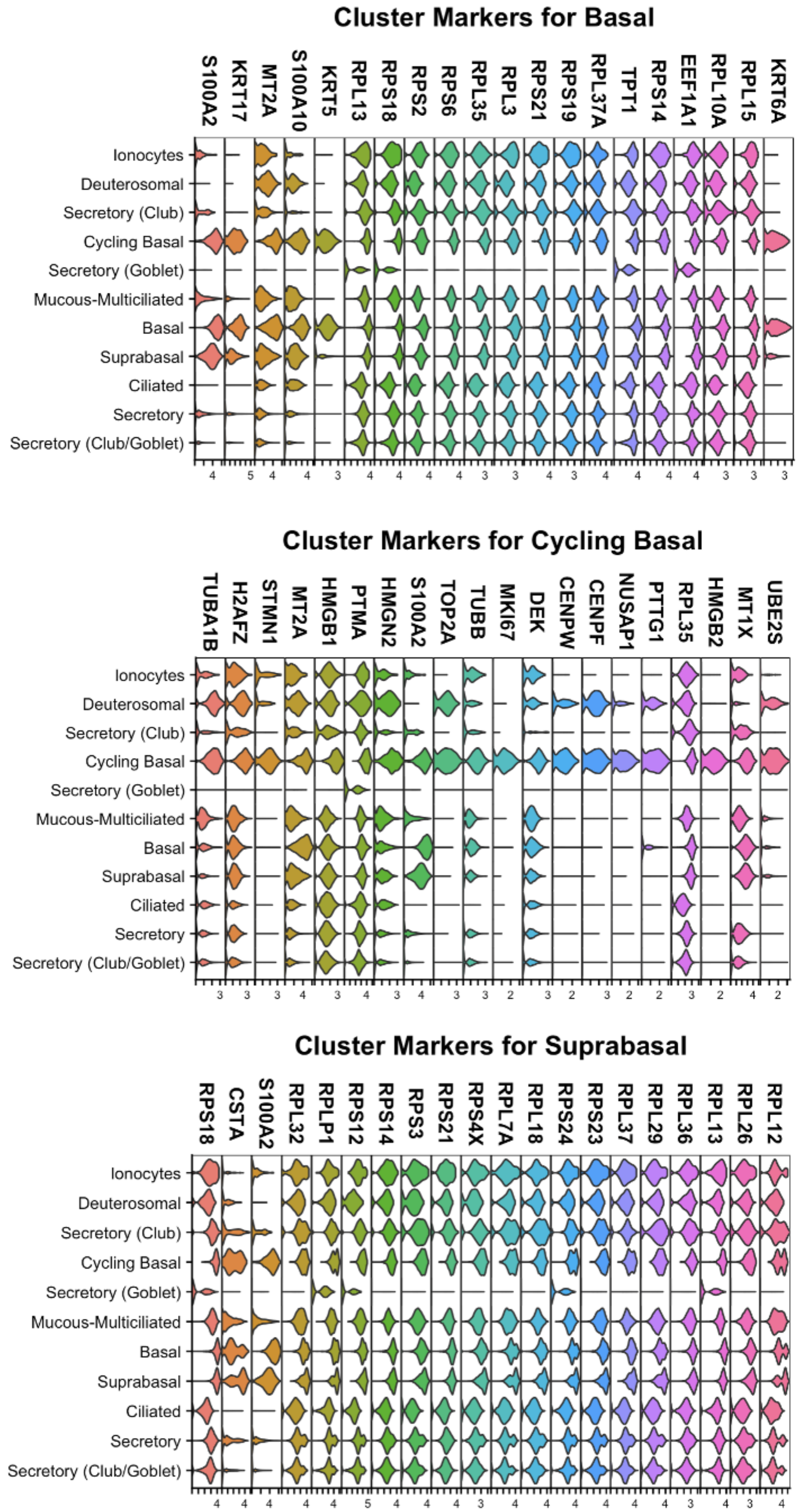

Cluster Markers for Basal
S100A2
KRT17
MT2A
S100A10
KRT5
RPL13
RPS18
RPS2
RPS6
RPL35
RPL3
RPS21
RPS19
RPL37A
TPT1
RPS14
EEF1A1
RPL10A
RPL15
KRT6A
Ionocytes
Deuterosomal
Secretory (Club)
Cycling Basal
Secretory (Goblet)
Mucous-Multiciliated
Basal
Suprabasal
Ciliated
Secretory
Secretory (Club/Goblet)
Cluster Markers for Cycling Basal
TUBA1B
H2AFZ
STMN1
MT2A
HMGB1
PTMA
HMGN2
S100A2
TOP2A
TUBB
MKI67
DEK
CENPW
CENPF
NUSAP1
PTTG1
RPL35
HMGB2
MT1X
UBE2S
Ionocytes
Deuterosomal
Secretory (Club)
Cycling Basal
Secretory (Goblet)
Mucous-Multiciliated
Basal
Suprabasal
Ciliated
Secretory
Secretory (Club/Goblet)
Cluster Markers for Suprabasal
RPS18
CSTA
S100A2
RPL32
RPLP1
RPS12
RPS14
RPS3
RPS21
RPS4X
RPL7A
RPL18
RPS24
RPS23
RPL37
RPL29
RPL36
RPL13
RPL26
RPL12
Ionocytes
Deuterosomal
Secretory (Club)
Cycling Basal
Secretory (Goblet)
Mucous-Multiciliated
Basal
Suprabasal
Ciliated
Secretory
Secretory (Club/Goblet)

C

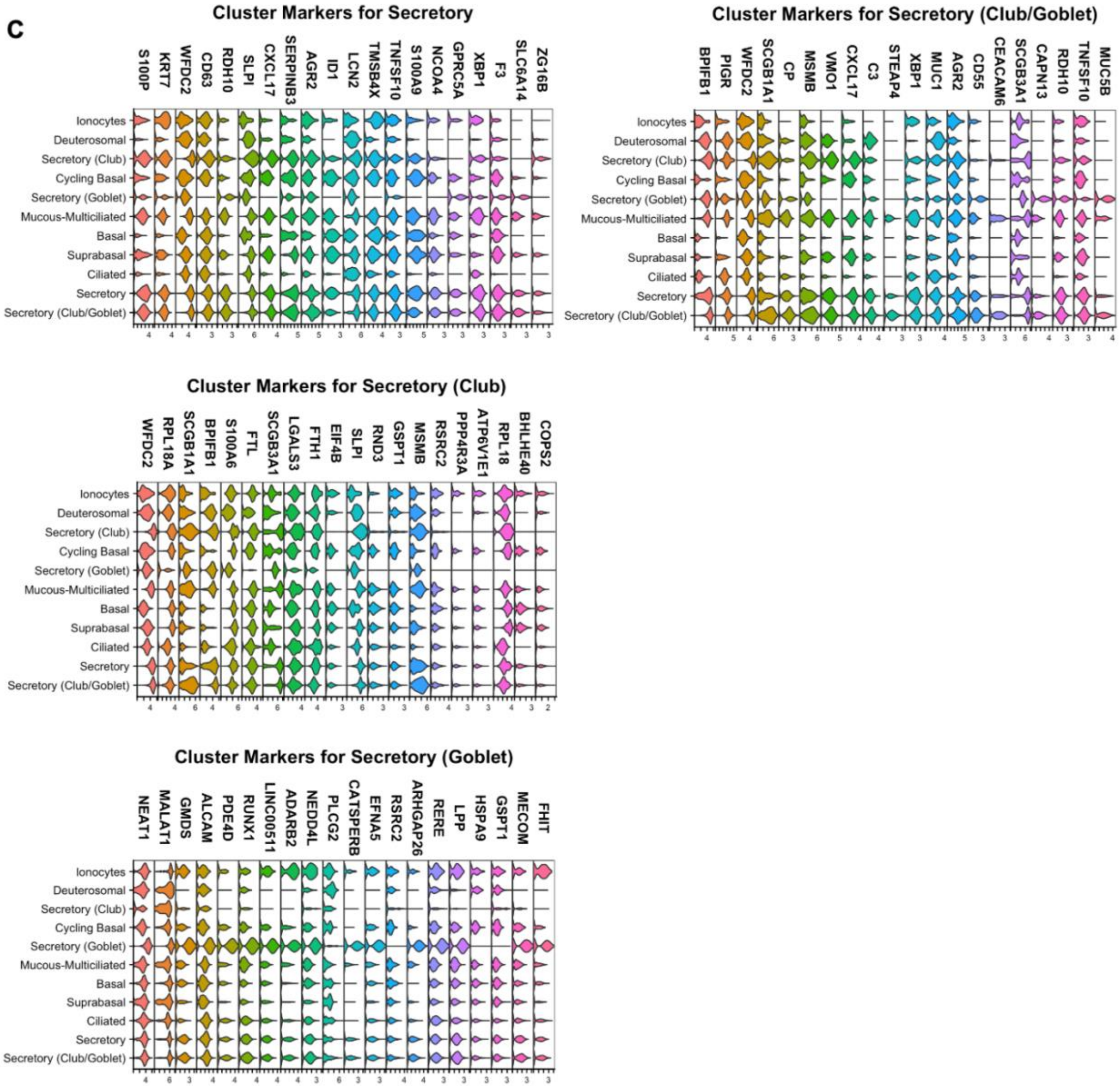
Cluster Markers for Secretory
S100P KRT7 WFDC2 CD63 RDH10 SLPI CXCL17 SERPINB3 AGR2 ID1 LCN2 TMSB4X TNFSF10 S100A9 NCOA4 GPRC5A XBP1 F3 SLC6A14 ZG16B
Cluster Markers for Secretory (Club/Goblet)
BPIFB1 PIGR WFDC2 SCGB1A1 CP MSMB VMO1 CXCL17 C3 STEAP4 XBP1 MUC1 AGR2 CD55 CEACAM6 SCGB3A1 CAPN13 RDH10 TNFSF10 MUC5B
Cluster Markers for Secretory (Club)
WFDC2 RPL18A SCGB1A1 BPIFB1 S100A6 FTL SCGB3A1 LGALS3 FTH1 EIF4B SLPI RND3 GSPT1 MSMB RSRC2 PPP4R3A ATP6V1E1 RPL18 BHLHE40 COPS2
Cluster Markers for Secretory (Goblet)
NEAT1 MALAT1 GMDS ALCAM PDE4D RUNX1 LINC00511 ADARB2 NEDD4L PLCG2 CATSPERB EFNA5 RSRC2 ARHGAP26 RERE LPP HSPA9 GSPT1 MECOM FHIT
Ionocytes
Deuterosomal
Secretory (Club)
Cycling Basal
Secretory (Goblet)
Mucous-Multiciliated
Basal
Suprabasal
Ciliated
Secretory
Secretory (Club/Goblet)

D

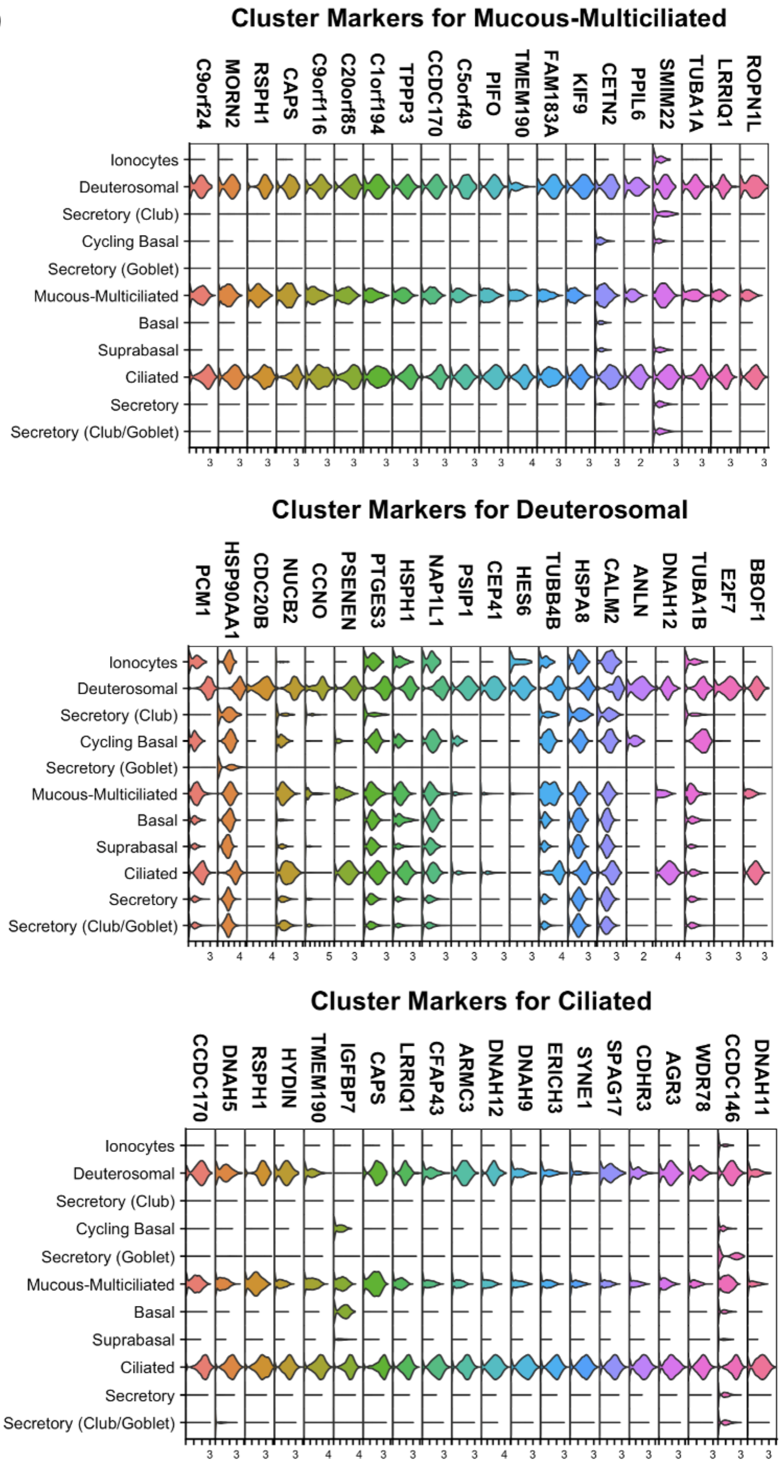


**Figure S3:** Violin plots visualizing top 20 differentially expressed genes for each secretory subtype. **(A)** Ionocytes. **(B)** Progenitor. **(C)** Secretory. **(D)** Ciliated.

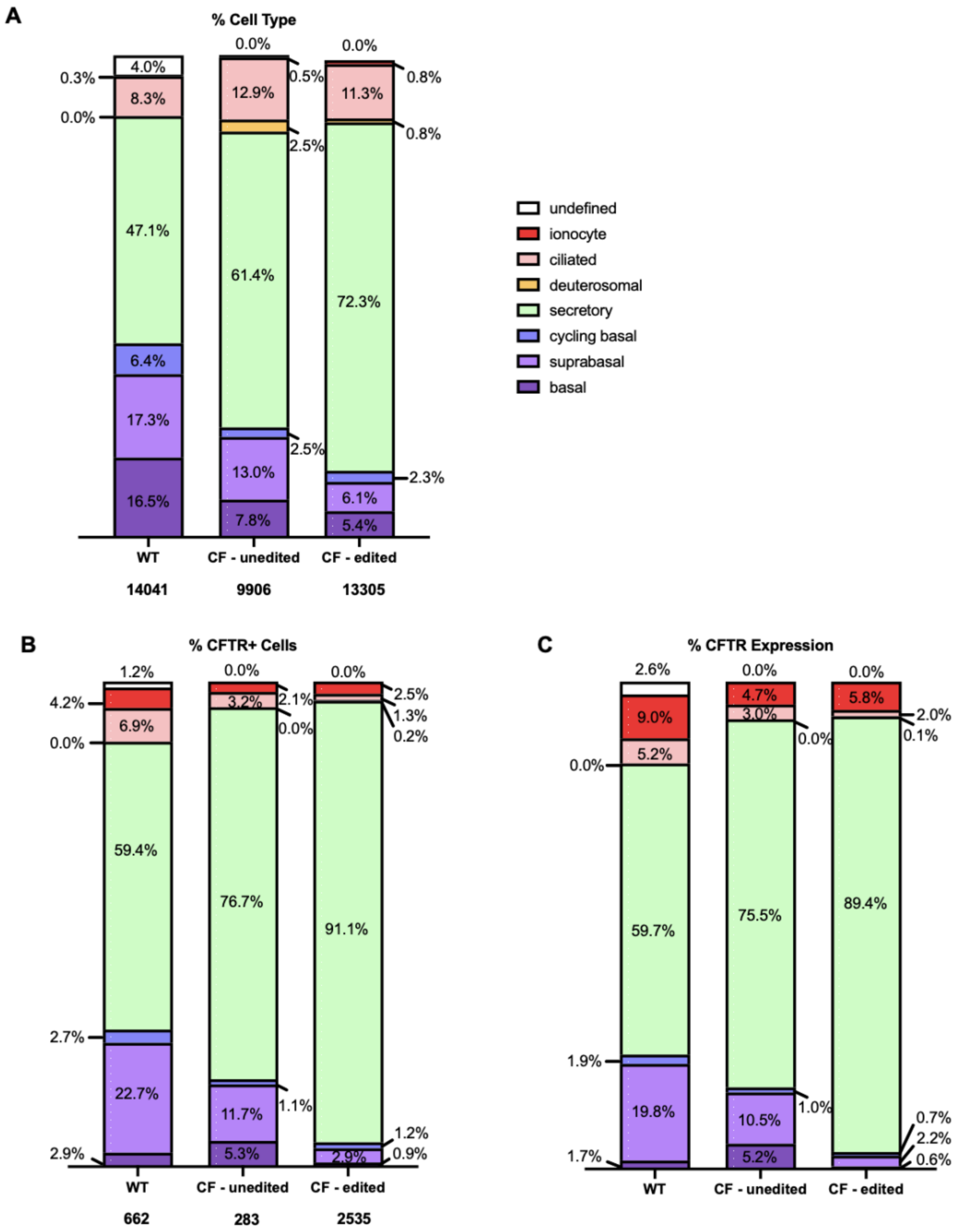


**Figure S4:** Graphical representations of cell type distribution, CFTR distribution and expression between WT (N=3), CF unedited (N=2) and CF edited (N=4) primary HNE samples differentiated for 21 days on ALI. Total number of cells per group is noted on the X-axis. Secretory is defined as combined secretory, secretory (club), secretory (goblet), secretory (club/goblet) and mucous-multiciliated populations. **(A)** Cellular composition between unedited and edited samples. **(B)** Percent CFTR+ cells across cellular subsets. **(C)** CFTR expression across cellular subsets, CFTR+ cells only.

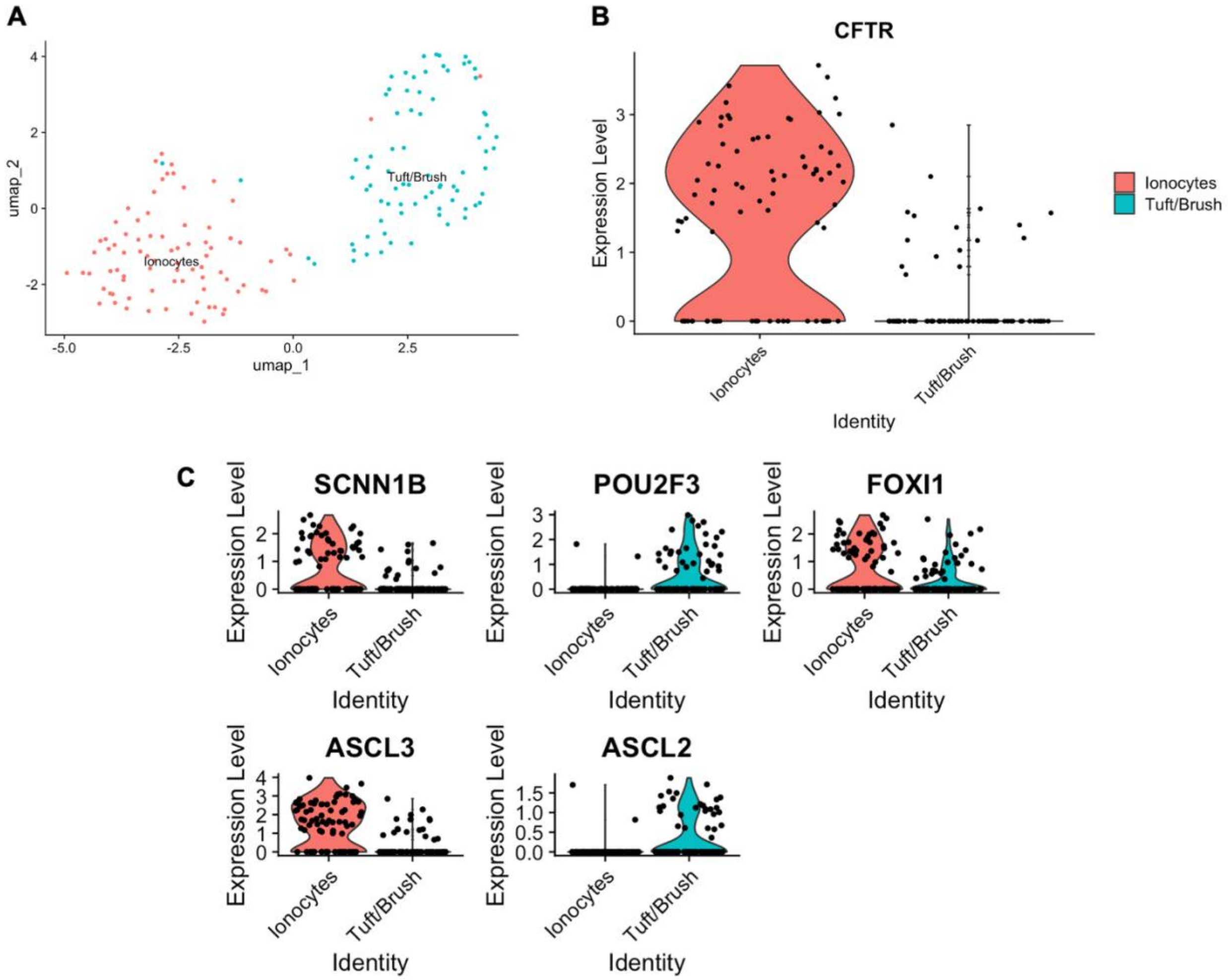


**Figure S5:** Subclustering of ionocytes reveals additional rare Tuft/Brush cell type. **(A)** Subclustering of ionocytes into ionocytes and tuft/brush cells. **(B)** CFTR Expression between the subclusters, visualized by Violin plot. **(C)** Expression levels of gene markers for ionocyte and tuft/brush cells, visualized by Violin plots.

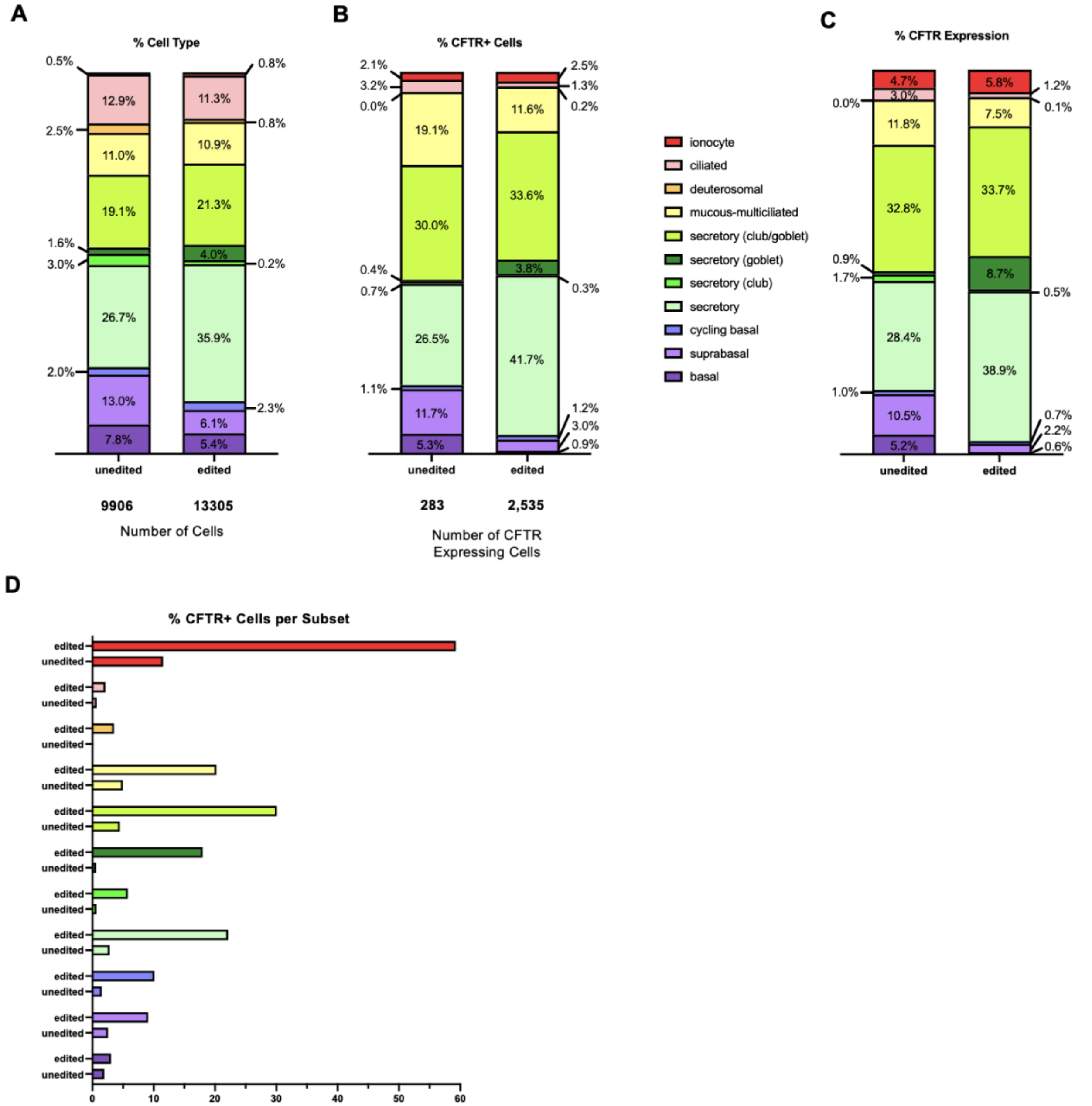


**Figure S6:** Graphical representations of cell type distribution, CFTR distribution and expression between unedited and combined edited primary HNE samples. **(A)** Cellular composition between unedited and edited samples. **(B)** Percent CFTR+ cells across cellular subsets. **(C)** CFTR expression across cellular subsets, CFTR+ cells only. **(D)** Percent CFTR+ cells contributable to either the unedited or edited samples, broken down by cellular subset.

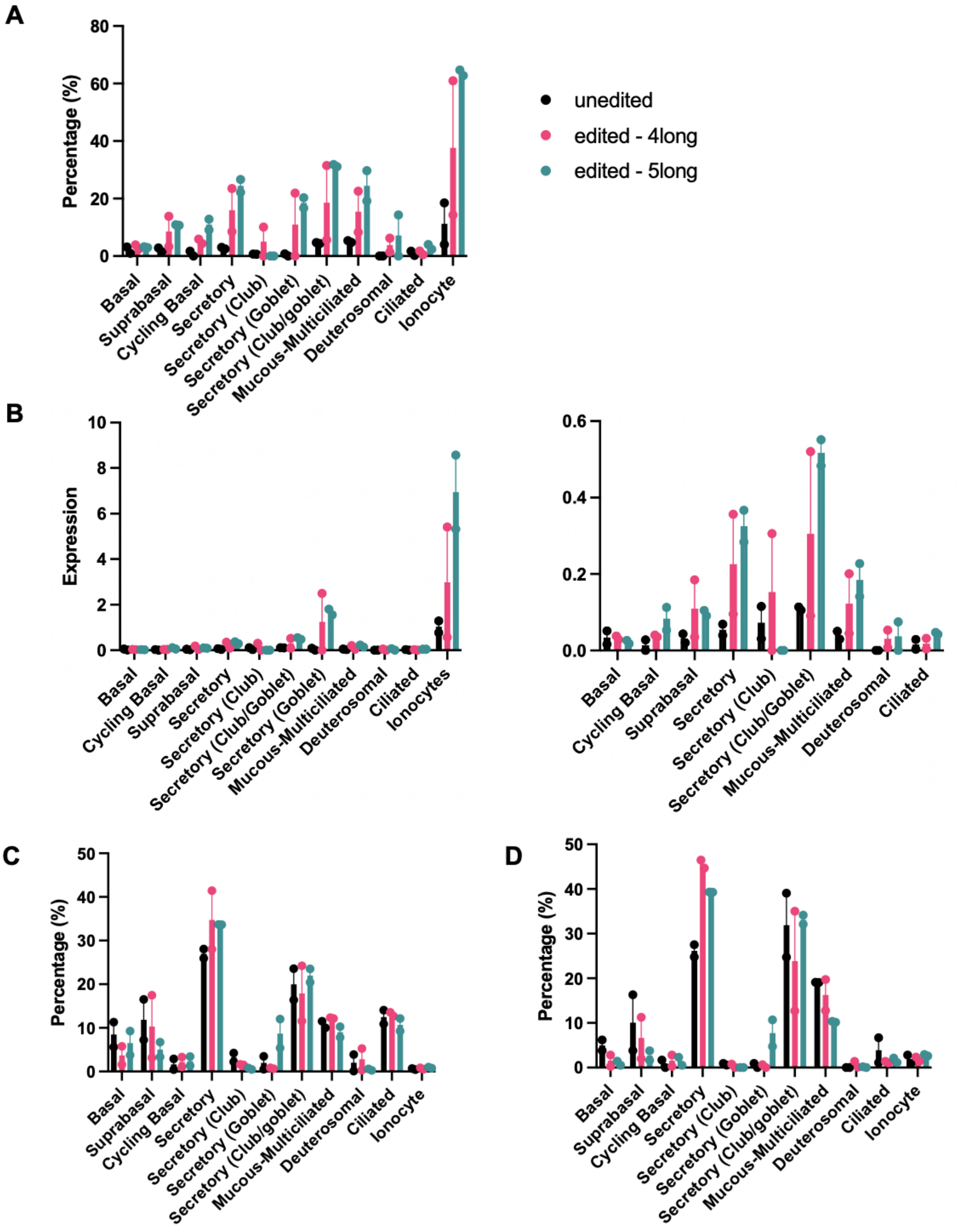


**Figure S7:** Individual values for **(A)** percent CFTR+ cells from each subset, **(B)** CFTR expression levels with (left) and without (right) the ionocyte subset, **(C)** percent cellular makeup, **(D)** percent CFTR+ cellular makeup. Data shown as mean ± SEM (N=2).

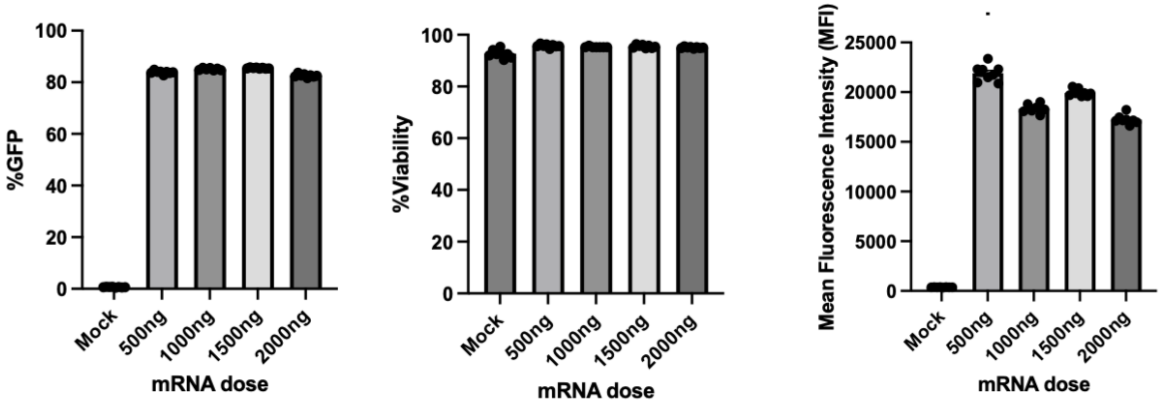


**Figure S8:** Flow cytometry results from nanoparticle transfection in immortalized CFBE cells bearing the 3120+1G>A variant with PBAE-E63: %GFP (left), %viability (middle), normalized mean fluorescence intensity (MFI) (right). Data shown as mean ± SEM (N=8).

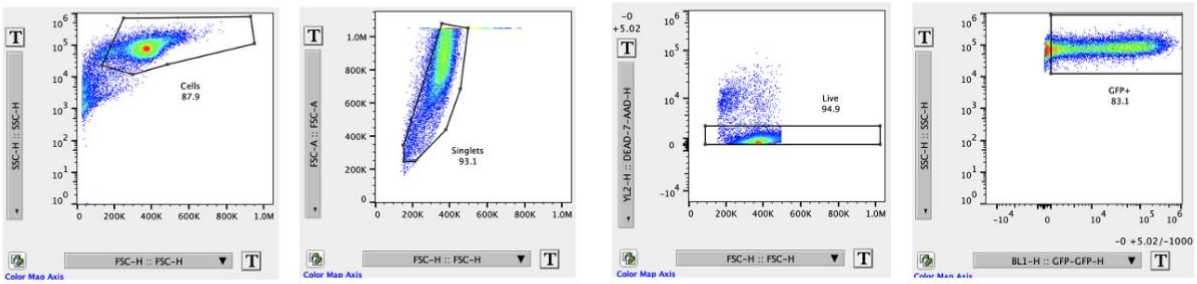


**Figure S9:** Flow cytometry gating for *in vitro* PBAE NP transfections in immortalized CFBE cells.

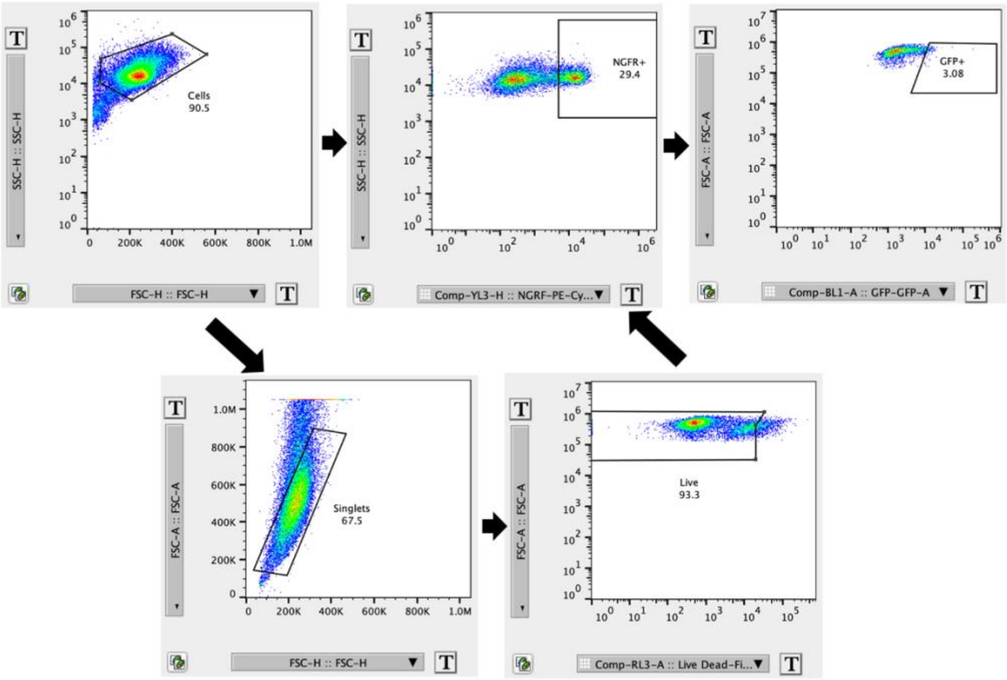


**Figure S10:** Flow cytometry gating strategy for primary HBE WT cells to identify specific lung cell population markers.

**Table S1: MiSeq gDNA sequencing primers for the 3120+1G>A variant.**

| Antigen | Color | Clone | Dilution | Supplier, Catalog No. |
|---|---|---|---|---|
| CD66c | R-phycoerythrin (PE) | 30-F11 | 1:600 | Biolegend; 103134 |
| PE/Cy7 | PE/Cy7 | M1/70 | 1:600 | Biolegend; 101217 |
| αTubulin | Alexa Fluor 405 | BM8 | 1:300 | Biolegend; 123128 |
| FcX | N/A | 93 | 1:100 | Biolegend; 101320 |

**Table S2: Comparing size, charge and PDI of PBAE NP[1] vs SORT LNP lung-targeting NPs[2]**

| | PBAE-E63 | SORT LNP 50% DOTAP | SORT LNP 100% DOTAP |
|---|---|---|---|
| **Size (nm)** | 157.8 | 113.1 | 118.2 |
| **Zeta Potential (mV)** | 34.45 | -0.52 | 25.50 |
| **PDI** | 0.14 | 0.22 | 0.20 |
| **w/w** | 30 | 40 | 40 |

**Table S3: Antibody list for primary HBE flow cytometry experiments.**

| Primer | Sequence (bold sequence is MiSeq tag) |
|---|---|
| Forward 5’ | **ACACTCTTTCCCTACACGACGCTCTTCCGATCTNNNN**GCTAATTCTTATTTGGGTTCTG |
| Reverse 5’ | **TGGAGTTCAGACGTGTGCTCTTCCGATCT**GCAATAGACAGGACTTCAACC |
